# Systematic microscopical analysis reveals obligate synergy between extracellular matrix components during *Bacillus subtilis* colony biofilm development

**DOI:** 10.1101/2022.06.16.496330

**Authors:** Michael Porter, Fordyce A. Davidson, Cait E. MacPhee, Nicola R. Stanley-Wall

**Affiliations:** Division of Molecular Microbiology, School of Life Sciences, University of Dundee, Dundee DD1 5EH, United Kingdom; Division of Mathematics, School of Science and Engineering, University of Dundee, Dundee DD1 4HN, United Kingdom; School of Physics and Astronomy, The University of Edinburgh, Edinburgh EH9 3FD, UK

**Keywords:** *Bacillus subtilis*, colony biofilm development, biofilm morphology, extracellular matrix, biofilm microscopy

## Abstract

Single-species bacterial colony biofilms often present recurring morphologies that are thought to be of benefit to the population of cells within and known to be dependent on the self-produced extracellular matrix. However, much remains unknown in terms of the developmental process at the single cell level. Here, we design and implement systematic time-lapse imaging and quantitative analyses of the growth of *Bacillus subtilis* colony biofilms. We follow development from the initial deposition of founding cells through to the formation of large-scale complex structures. Using the model biofilm strain NCIB 3610, we examine the movement dynamics of the growing biomass and compare with those displayed by a suite of otherwise isogenic matrix-mutant strains. Correspondingly, we assess the impact of an incomplete matrix on biofilm morphologies and sessile growth rate. Our results indicate that radial expansion of colony biofilms results from division of bacteria at the biofilm periphery rather than being driven by swelling due to fluid intake. Moreover, we show that lack of exopolysaccharide production has a negative impact on cell division rate, and the extracellular matrix components act synergistically to give the biomass the structural strength to produce aerial protrusions and agar substrate-deforming ability.

## Introduction

Bacteria growing at an interface between two phases often form biofilms. In the laboratory, biofilms can present in many different forms including floating on the surface of a stationary liquid (pellicle biofilm), within a liquid but adherent to an object (submerged biofilm) or on a solid substrate open to air (colony biofilm) (O’Toole et al., 2000). Common across each form of biofilm is a self-produced extracellular matrix comprising polymeric proteins and saccharides, extracellular DNA, surfactants, and other substances (Flemming and Wingender, 2010). The biofilm matrix provides cohesion within the biomass, protects cells from environmental and chemical assault, and aids in hydration and nutrient acquisition (Dragoš and Kovács, 2017). The matrix also confers “emergent” properties to the collective (Flemming et al., 2016).

A key feature of laboratory-grown colony biofilms is the complex architecture that develops. Patterns of ridges and wrinkles are often seen, where the ordered and reproducible nature of the resulting architecture can be indicative of the species under investigation (Gordon et al., 2019). These complex patterns in the growing colony biofilm biomass are a response to the physical environment (for example, stiffness of the substrate), the availability of both nutrients and oxygen, and are dependent on the composition of the biofilm matrix (Fei et al., 2020).

Microscopical analyses of localised areas of colony biofilms have been performed to observe the physical movement of Gram-negative bacteria forming colony biofilms. *Pseudomonas aeruginosa*, in the context of a colony biofilm, divide into discrete cells and use the Type IV pilus to move across the surface and expand the biomass at the very edge of the community (Meacock et al., 2020). At higher cell densities further into the biomass, varying velocities of this pilus-driven movement generate degrees of orderly packing of cells, which has an impact on the overall biofilm morphology (Meacock et al., 2020). Additionally, a model has been proposed to simulate the growth and morphology-generating forces of *Vibrio cholerae* biofilms, accounting for the friction of cells on the substrate, the presence of extracellular matrix and the diffusion of nutrients (Fei et al., 2020).

*Bacillus subtilis* is a Gram-positive soil-dwelling bacterium that is frequently used to study biofilms (Arnaouteli et al., 2021). While recent systematic analyses of pellicle biofilm formation have been conducted and revealed complex dynamics in cell behaviour (Krajnc et al., 2022; Sanchez-Vizuete et al., 2022), the same has not been done for biofilms growing on a solid substrate. In contrast to *P. aeruginosa* or *V. cholerae*, *B. subtilis* grows on a semi-solid substrate as long chains of connected cells (Arnaouteli et al., 2019; O’Toole and Kolter, 1998; Yan et al., 2016). Work to elucidate the forces at play regarding movement of these chains of cells has been undertaken. As the cells grow on an agar surface and form microcolonies, deformations in the orderly chains occur at higher cell densities (Yaman et al., 2019), with contributing structural effects from various matrix molecules (Seminara et al., 2012; Cámara-Almirón et al., 2020; Dragoš et al., 2017).

It is well documented that colony biofilms formed by bacterial strains lacking genes that lead to matrix production display morphologies of reduced complexity (Branda et al., 2001). Multiple end-point techniques have been employed to probe the importance of the matrix to biofilm architecture and development such as: macro-scale microscopy (Branda et al., 2001); destructive techniques including thin-section preparation for light and electron microscopy (Hobley et al., 2013; Watnick et al., 2001); or biofilm disruption to single cells for flow cytometry (Veening et al., 2006). While these approaches are informative for questions of protein localisation within the biomass or for a snapshot of morphology in the presence or absence of specific matrix components, we lack a motion-based non-destructive analysis of the cell growth and movement responsible for the emergence of structural complexity in the mature community.

Here we present a systematic time-lapse microscopy-based examination of colony biofilm growth by *B. subtilis* NCIB 3610. We cover defined periods of growth including the point of initial deposition of the founding cells to the agar surface, through to maturation of the colony biofilm architecture. We attain an unprecedented understanding of the different stages of colony biofilm formation and ascribe quantitative phenotypic differences to genetic mutations altering the constituents of the matrix. We reveal the contribution of the polymers of the matrix to morphological development of biofilms by analysing the behaviour of the tissue and link the matrix components with the overall growth mechanics of the *B. subtilis* colony biofilm.

## Results

### Gross colony biofilm morphology varies with matrix composition

Through macro-scale imaging of colony biofilms, the inability of *B. subtilis* to produce the full array of biofilm matrix components has been linked with *i)* gross morphological differences in colony biofilm architecture (Branda et al., 2001), *ii)* the footprint occupied by the biomass (Kesel et al., 2017), and *iii)* the community’s ability to deform the substratum on which it is grown (Zhang et al., 2015). Here, reflected light images of colony biofilms formed by NCIB 3610, and a suite of otherwise isogenic mutant strains carrying deletions in the *tapA*, *sipW*, *tasA*, *tapA tasA*, *epsH*, and *bslA* coding regions, demonstrate these impacts (Fig. 1A) (see Strain Table). Qualitatively, as expected, each of the deletion strains showed a gross reduction in structural complexity as compared to the parental strain colony biofilm (e.g., Branda et al., 2001; Romero et al., 2010), although it should be noted that the exact colony biofilm morphology varies with each genotype. The mutant strains also showed differences in the area occupied by their colony biofilms (Fig. 1B). Specifically, at 24 hours of growth, the footprints of the *epsH* and *tapA tasA* mutant colonies were smaller than those formed by NCIB 3610 (p<0.0001), and by 48 hours all the mutant strains occupied significantly smaller footprints (Fig. 1B).

**Figure 1.**
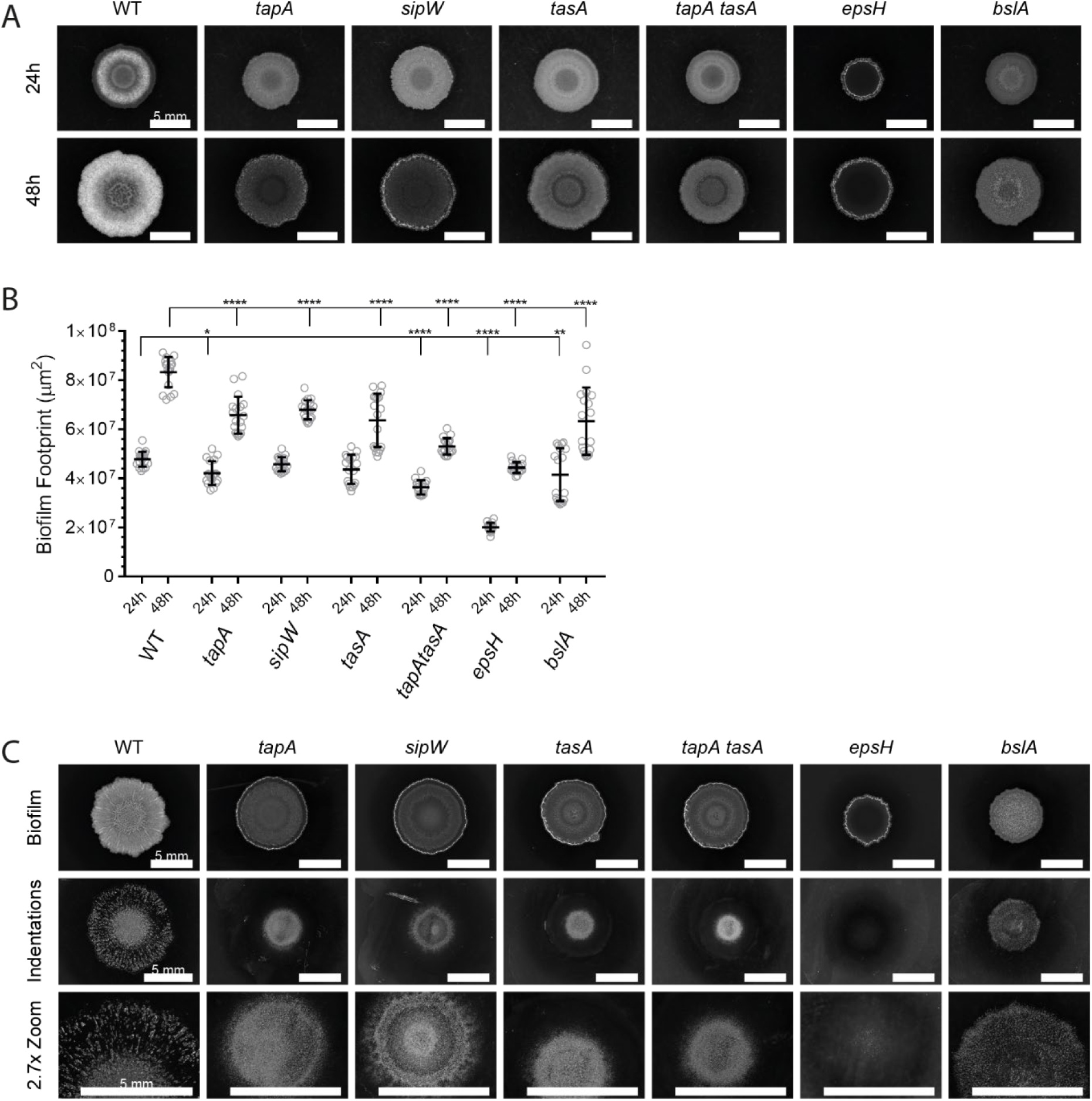
Removal of each *B. subtilis* matrix molecule has a distinct impact on colony biofilm formation. **(A)** A macro-scale view of colony biofilms of NCIB 3610 (WT) and derived strains carrying specific deletion mutations for matrix component genes (*tapA* NRS3936, *sipW* NRS5488, *tasA* NRS5267, *tapA tasA* NRS5748, *epsH* NRS5906, *bslA* NRS2097). The same biofilm is presented at 24h and 48h for each strain as indicated, incubated at 30°C; **(B)** Footprints of colony biofilms formed by the indicated strains at 24h and 48h after cell deposition. Reflected light macroscopic images were segmented in Matlab and areas calculated from the pixel size of the images. n=18. *p = 0.0106, **p = 0.0033, ****p < 0.0001; **(C)** Indentation patterns caused by colony biofilms grown at 30°C for 72 hours revealed by removal of the biomass. Scale bars represent 5 mm.

Colony biofilms formed by NCIB 3610 force indentations to develop in the agar substratum, thought to be caused by osmotic pressure exerted by the presence of EPS in the biofilm (Zhang et al., 2015). As expected, multiple indentations in the agar under NCIB 3610 colony biofilms could be seen below the initial deposit of founding cells (within the ‘coffee ring’ of high-density cells produced by inoculum evaporation), with further indentations in the agar extending in a radial pattern to the colony biofilm periphery (Fig. 1C). Moreover, as previously reported only low levels of substratum deformation could be induced by the *epsH* strain (Fig. 1C) (Zhang et al., 2015). Our new analysis of the other matrix deletion strains revealed that the only matrix mutant to show a peripheral pattern of substratum indentation was the *bslA* strain, although visually the indentations formed appeared to be of lower frequency and less pronounced than those formed by NCIB 3610. The remaining matrix mutant strains (*sipW, tapA, tasA, tapA tasA*) deformed the substratum to varying degrees in the agar directly under the initial inoculum, but deformations were mainly absent under the colony biofilm periphery. We therefore conclude that absence of any of the matrix components alters how the biofilm interacts with the substratum.

### Growth rate in the initial colonisation phase is not linked with the mature biofilm footprint

We proposed that novel insight to the microscale processes that influence the macroscale presentation of colony biofilms may be gained by in-depth systematic imaging that quantifies the dynamics of cell movement during the formation process. Therefore, a series of confocal imaging experiments were designed and conducted (Fig. S1, Fig. S2B). The initial experiments involved collecting data from the earliest point of founder cell deposition onto the agar substrate (0-10 hours) (Fig. S1A). The starting planktonic NCIB 3610 cultures were normalised to an optical density of 1 at OD_600_ and a 1 µl inoculum was spotted onto the agar surface. The founding cells contained a ratio of 20% GFP-positive and 80% GFP-negative cells to aid visualisation, by adding contrast to the biomass that quickly becomes optically dense. The inoculum was dried onto the surface of the agar forming a patch of founding cells in a characteristic ’coffee ring’ pattern (Wang et al., 2016) (Fig. Movie 1). Over time, the bacteria increased in number and started to grow as chains (Fig. 2A), forming a coherent biomass where the cells filled the agar surface within the coffee-ring (Fig. Movie 1).

**Figure 2.**
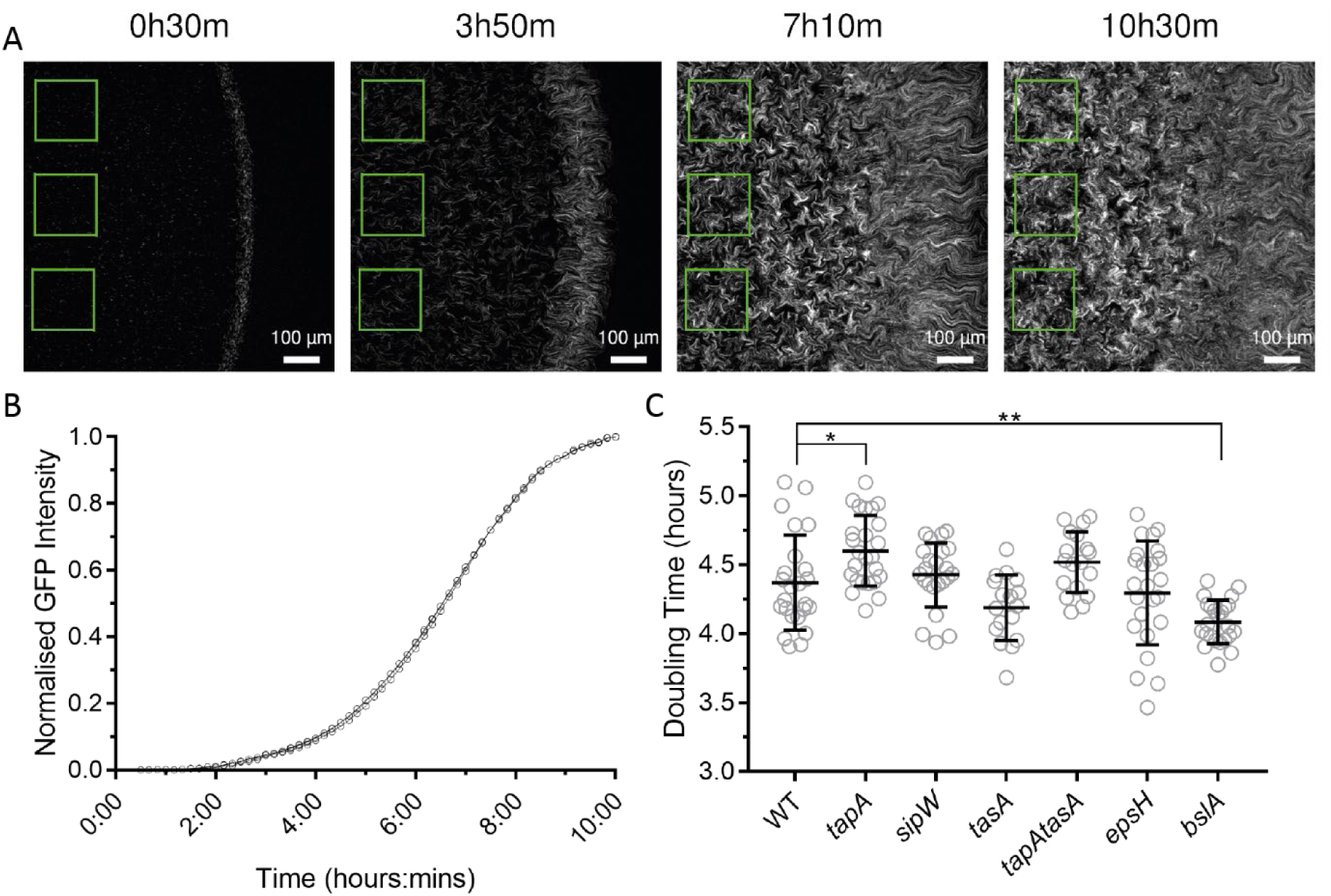
Quantitation of initial cell growth. **(A)** Example analysis process where static ROIs (green squares) are placed towards the centre of biomass and used to read the fluorescence signal generated by the GFP-positive cells over a 10-hour period. For experimental setup see Fig. S1A; **(B)** Representative example of fluorescence signals from three ROIs normalised to their maximum value in a single time-lapse series of NCIB 3610/NRS1473; **(C)** Calculated doubling times inferred from the maximum gradient of the logistic GFP fluorescence graphs (B) for the indicated strains over the first 10.5 hours of colony biofilm growth (WT NCIB 3610/NRS1473, *tapA* NRS3936/NRS6723, *sipW* NRS5488/NRS6718, *tasA* NRS5267/NRS6724, *tapA tasA* NRS5748/NRS6727, *epsH* NRS5906/NRS6728, *bslA* NRS2097/NRS5131). Individual data-points are shown with over-laid bars representing the mean and standard deviation for each population of data. n = 17-25. *p = 0.0181, **p = 0.0023.

We systematically assessed the behaviour of the suite of biofilm matrix mutants over the same timeframe (see Strain Table) (Fig. Movie S1 A – F). As with NCIB 3610, each of the matrix mutants contained populations of cells that were physically constrained within a coffee-ring formed upon deposition of the inoculum (Fig. 1A). To determine if differences in mature colony biofilm morphology and footprint were due to changes in growth rate during colonisation of the agar surface, the normalised mean intensity of GFP signal was measured every 10 minutes over a 10-hour period, with the GFP intensity value serving as a proxy for cell density measurements (see materials and methods) (Fig. 2A) (Fig. 2B) (Fig. 2C). There was a difference in the mean doubling times for each of the matrix mutant strains compared to NCIB 3610, with the *tapA* strain being slightly slower and *bslA* strain slightly faster (p=0.018 and p=0.002 respectively, Fig. 2C). Although statistically different, most of the calculated doubling times for these two strains were values within the standard deviation of NCIB 3610 (Fig. 2C,). Therefore, it is unlikely that differences in the ability of the biofilm matrix mutant strains to initially colonise the agar surface of the inoculation zone would lead to consistently smaller colony biofilm footprint or architecture.

### Biofilm edge expansion is linked with biofilm matrix molecules

Concurrent with growth of the cell population that was constrained within the central coffee-ring zone, chains of NCIB 3610 cells emanated from the exterior boundary of the coffee-ring and expanded radially across the agar surface increasing the footprint (Fig. Movie 1) (Arnaouteli et al., 2019). To record expansion of the colony biofilm from this point forward, a separate imaging experiment following growth of the monolayer cell chains was initiated. At 10.5 hours after the cells were initially deposited, imaging data were collected for the next 12-15 hours. (Fig. S1B) (Fig. Movie 2, Fig. 3A). By comparing images collected over time we observed that the NCIB 3610 chains travelled >700 µm radially across the agar before leaving the field of imaging. The chains typically traversed the field of view at a rate of 4.25 µm/minute and with complex motion of the constituent loops and chains in the expanding biomass (Fig. Movie 2). We also examined the biofilm matrix mutant strains during this phase of colony biofilm formation (Fig. Movie S2 A - F). All the strains grew chains of cells emanating from the coffee ring biomass and we calculated the rate of expansion of the biomass across the agar surface for each genotype (see materials and methods). All bar one of the matrix mutant strains displayed a significantly slower mean expansion rate than NCIB 3610, ranging from 0.74 µm/minute for the *epsH* strain (p<0.0001) to 2.54 µm/minute for the *bslA* mutant strain (p=0.001). The *sipW* strain mean expansion rate was not significantly different from NCIB 3610 at 3.72 µm/minute, but the data showed two populations – some at the upper bounds of NCIB 3610 rates and the others as slow as the *tapA* and *bslA* strains (Fig. 3C). Overall, these data link specific biofilm matrix molecules to expansion of the biomass from the coffee-ring and are consistent with observed differences in the final colony biofilm footprint (Fig. 1A, 1B).

**Figure 3.**
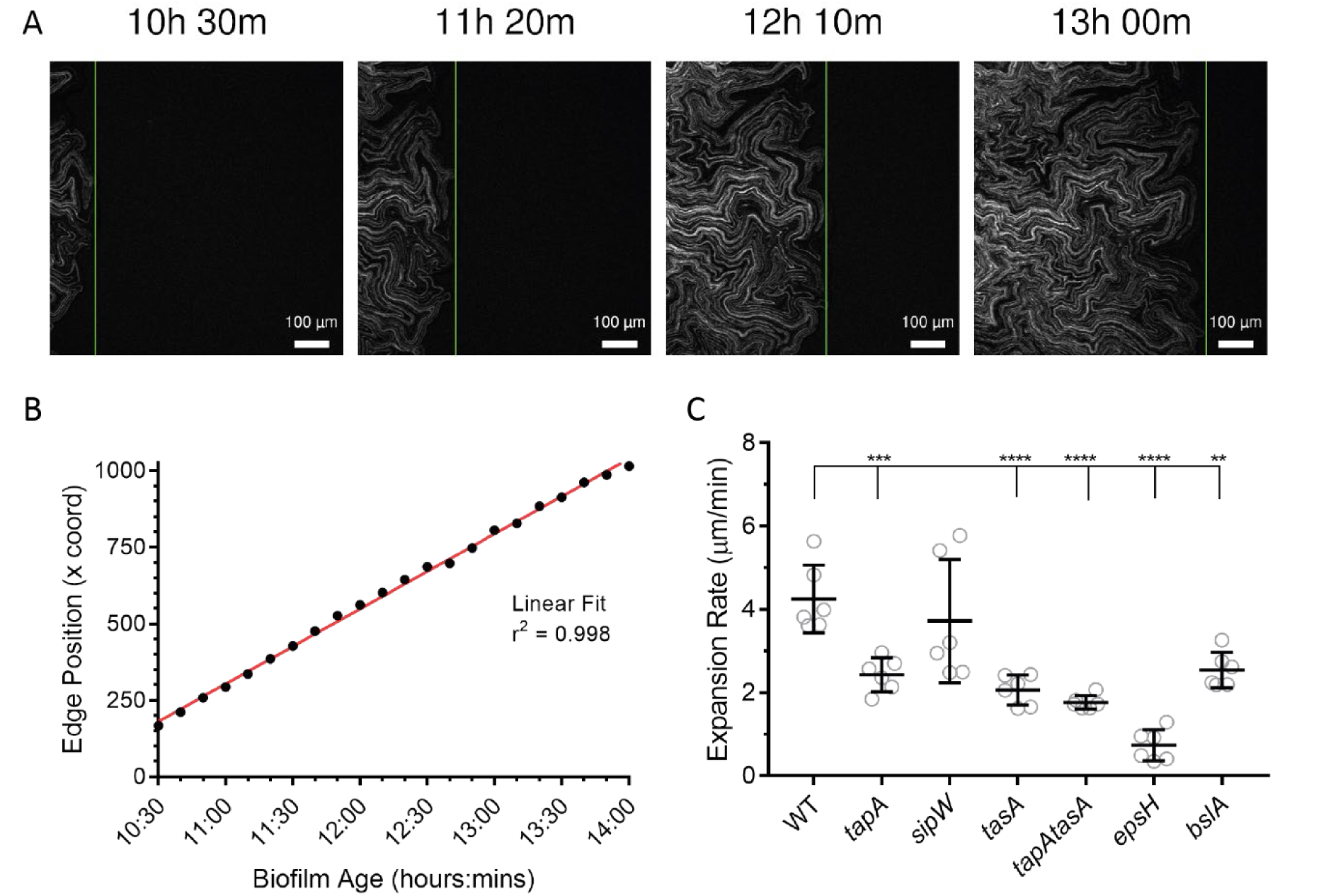
Expansion of cell chains from the biofilm periphery. **(A)** Example time points of time-lapse imaging of an expanding NCIB 3610/NRS1473 colony biofilm. The time stamp on each panel gives the approximate age of the colony biofilm based on the point when cells were deposited onto the agar. The vertical green line denotes the automatically detected segmented edge of the colony biofilm, the position of which was recorded. For experimental setup see Fig. S1B; **(B)** Example plot of the position of the outside edge of an NCIB 3610 biofilm over time across the image space (black circles). A linear fit can be applied to these data with high confidence (red line). The linear fit was used to calculate expansion rate of the cell chains across the surface; **(C)** Expansion rates calculated from fitted lines exemplified in (B). Individual data points are shown, overlaid with bars representing the mean and standard deviation of each population (WT NCIB 3610/NRS1473, *tapA* NRS3936/NRS6723, *sipW* NRS5488/NRS6718, *tasA* NRS5267/NRS6724, *tapA tasA* NRS5748/NRS6727, *epsH* NRS5906/NRS6728, *bslA* NRS2097/NRS5131). N=6. **p = 0.001, ***p = 0.005, ****p < 0.0001.

### The absence of the biofilm exopolysaccharide increases cell cycle times

Slower expansion across the agar surface by chains of cells produced by the biofilm matrix mutants could be the consequence of slower cell division rates at the single cell level culminating in reduced biomass production. Therefore, to calculate the time taken for a single cell to divide during biofilm expansion, strains were constructed in NCIB 3610 and matrix-gene deletion backgrounds that harboured a gene encoding an FtsZ-GFP fusion protein inserted at the native *ftsZ* locus. The strains also constitutively expressed *mKate2* (Strain Table). FtsZ is one of the first proteins to localise at the mid-point of dividing cells, forming a scaffold (‘z-ring’) for the machinery that will carry out cytokinesis (Fig. Movie 3) (Gamba et al., 2009). After cytokinesis the z-ring disassembles are re-forms at the mid-point of the daughter cells. The interval between the formation of z-rings of parent and daughter cells can therefore be monitored to calculate the generation time (Fig. 4B). The presence of the FtsZ-GFP fusion protein did not impact growth, as no phenotypic difference in the colony biofilm morphology compared with the parental strain lacking the reporter fusion were detected (Fig. S2A).

**Figure 4.**
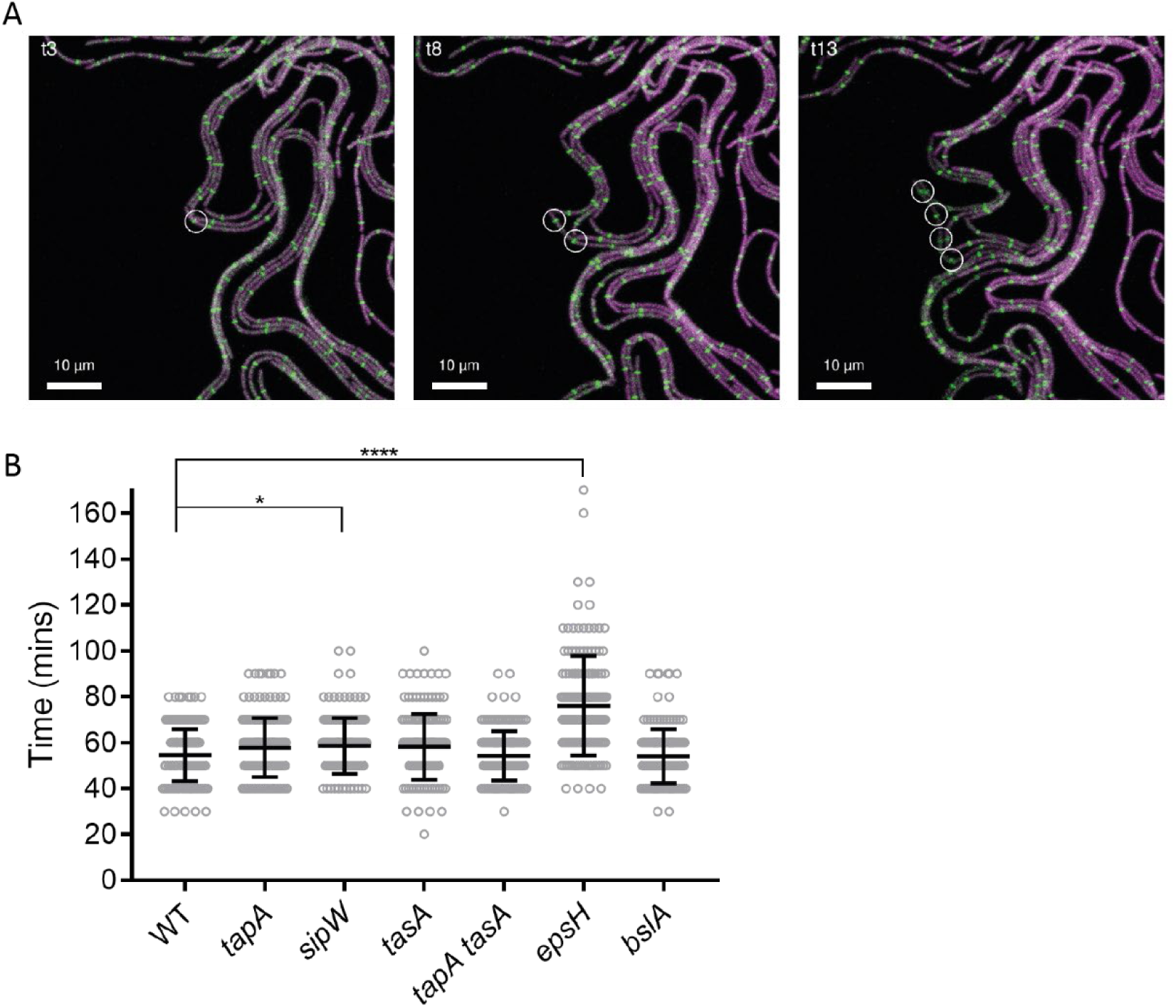
Single cell division time within the context of a colony biofilm. **(A)** Exemplar of CLSM time-lapse imaging of FtsZ-GFP (green) and mKate2 (magenta) expressing cells (NRS6781) mixed at a 1 in 5 starting ratio with NCIB 3610. A nascent Z-ring is marked at time-point 3 (circle, panel 1) and followed to observe the appearance of daughter cell Z-rings at time-point 8 (panel 2) and granddaughter Z-rings at time-point 13 (panel 3). The interval between time-points is 10 minutes, allowing calculation of the duration of division events. For experimental setup see Fig. S2B; **(B)** Cell division times for NCIB 3610 (WT) and matrix gene mutant strains as indicated. n=152 minimum division events. Black bars show mean and standard deviation of the data (WT NCIB 3610/NRS6781, *tapA* NRS3936/NRS6789, *sipW* NRS5488/NRS6794, *tasA* NRS5267/NRS6795, *tapA tasA* NRS5748/NRS7525, *epsH* NRS5906/NRS7526, *bslA* NRS2097/NRS7527). *p = 0.0401, ****p < 0.0001

To measure the generation time at the single cell level in conditions where the biofilm matrix is produced, the suite of strains was grown under biofilm-forming conditions for 12 hours prior to high resolution imaging for multiple generations (Fig. 4A, Fig. S2B). These analyses provided data to calculate the mean generation time for each strain growing on a solid surface (Fig. 4B). NCIB 3610 showed a distribution of doubling times, with a mean of 54 minutes (±11 minutes SD) (Fig. 4B). Analysis of the biofilm matrix mutants yielded comparable values (Fig. 4B), except for the *sipW* strain (p = 0.04), with a slight increase in mean division time of four minutes compared to NCIB 3610 but with similar standard deviation, and more significantly the *epsH* strain took 79 minutes (±22 minutes SD) to divide (Fig 4B). These findings reveal that the absence of exopolysaccharide during sessile growth impacts cell division, and this is likely to contribute to the reduction of the footprint of the *epsH* mutant compared to NCIB 3610.

### A reduction in collective biomass movement is associated with an incomplete matrix

Alongside differences in the expansion rate attained by the biofilm matrix mutants, variations in the form of motion displayed by the cell chains expanding across the agar surface were observed. In NCIB 3610, the density of the biomass to the inside of the expanding edge of the colony biofilm increased to the point where lateral movement became restricted. At this point, the chains of cells apparent in the biomass (due to the definition afforded by the GFP-positive cells contrasting with the GFP-negative) became bundled and appeared twisted, ultimately leading to slower moving biomass above the plane of the agar (Fig. Movie 4, arrows) (Yaman et al., 2019). The formation of vertical structures in the peripheral zone of the biomass is consistent with the behaviour of cells within the coffee-ring zone which also appear rise because of physical constraint (Fig. Movie 1). Likewise, at this point, as the cells pile up vertically, they continue to increase the overall biomass. In contrast to NCIB 3610, the expanding chains of the matrix mutant strains, except for the *bslA* mutant, formed fewer 3-dimensional structures and maintained a monolayer appearance in a larger zone of the colony biofilm periphery (Fig. S3 WT vs *sipW*). These observations are consistent with the static single time point perspective of the biomass obtained by confocal imaging of later stage colony biofilms at both the periphery and interior (Fig. Movie S2 A - F, Fig. 9).

To compare the dynamics of motion across the array of strains, we marked and tracked the 2D movement of 36 visibly distinct features in the biomasses (six features were followed per image, six images were in each dataset, and the features selected were distributed across the initial biomass in the field of view imaged) (see Materials and Methods) (Fig. Movie S3). The field of view imaged is positioned at the edge of each biofilm such that movement in a straight line along the x-axis from left to right is equivalent to, in the context of a polar coordinate system of a whole biofilm, increasing in radius without deviating in angle theta from the centre (radial growth). We analysed the path, rate of motion, and change in distance from the biofilm edge for each feature for the maximum duration of the fastest-moving features, to the point at which the edge of the biofilm left the field of view (which was one hour and 40 minutes). Plotting of feature trajectories from NCIB 3610 biofilms revealed the direction of movement is not strictly aligned with the x-axis, with features deviating to different degrees along the y-axis (Fig. S4). To compare the types of tracks of each genotype, the starting point for each trajectory was normalised to the origin and overlaid in separate plots (Fig. 5A). Comparing all the feature trajectories of the matrix mutant strains to those of NCIB 3610 revealed differences in the motion profiles - for example, the path taken by features within the *sipW* strain colony biofilm tended to extend outwards along the x-axis to a similar degree to, but less prone to motion along the y-axis than, those of NCIB 3610 (Fig. 5A *sipW*). Isolating the x-axis component of these trajectories revealed the *sipW* strain was the only mutant strain that did not produce feature tracks that were significantly shorter than NCIB 3610 in this direction (Fig. 5B, p=0.1617 compared to p<0.0001 for all other mutant strains). Isolating the y-axis component of the trajectories shows that all the mutant strains had a significantly reduced range of motion away from an x-axis trajectory compared to NCIB 3610 (Fig. 5C, p=0.0008 for *tapA* mutant strain, all other mutant strains p<0.0001).

**Figure 5.**
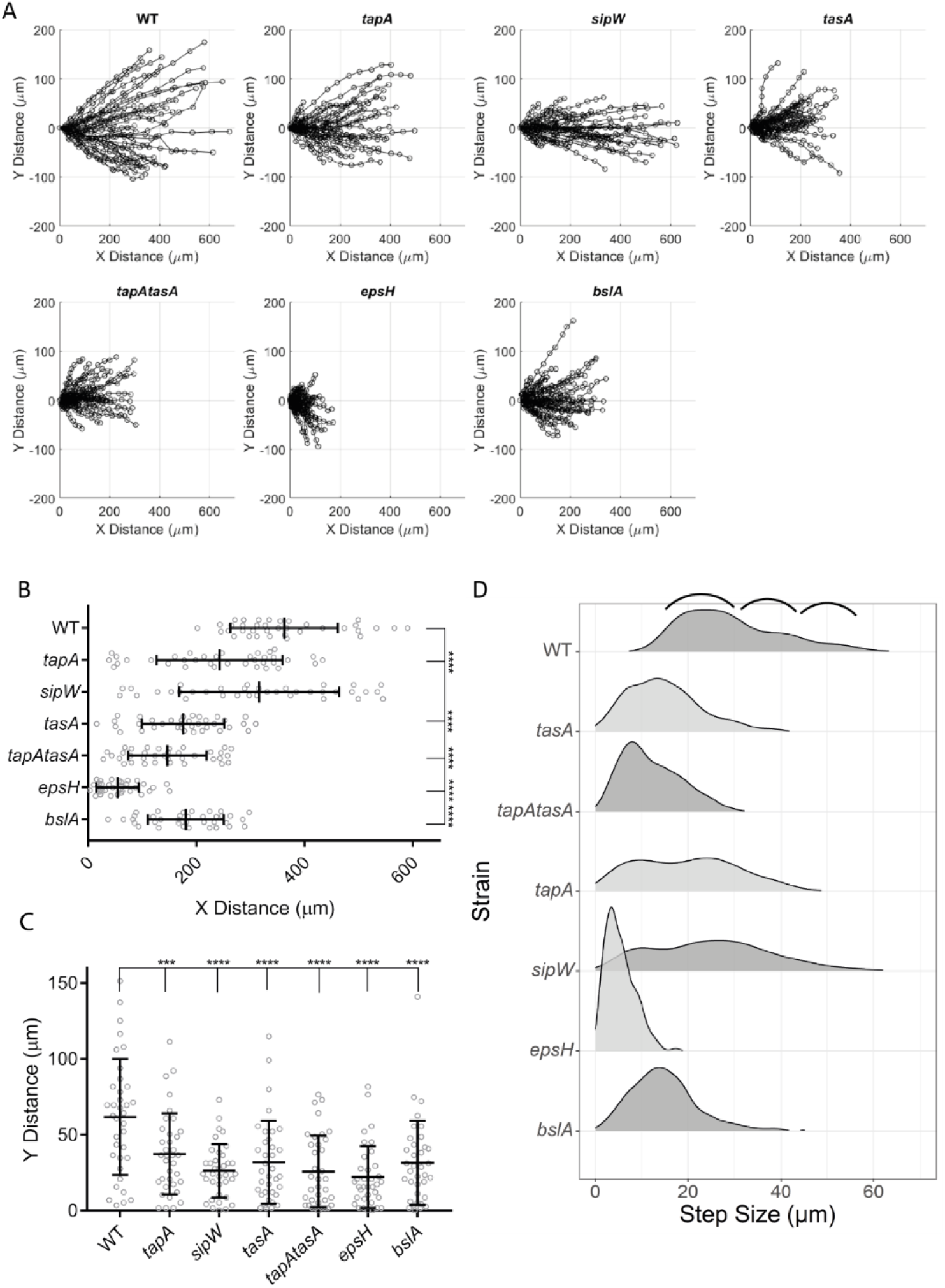
Biomass landmark motion analysis. **(A)** Tracks of landmarks represented in (Fig. S4) with their starting positions normalised to position (0,0) to display, for the strains indicated, the range of motion of biomass in the radial direction of biofilm growth (x-axis) and the range of deviation from this direction (y-axis); **(B** and **C)** Extraction of individual components of landmark trajectories motion from (A); **(B)** The X component is the range of motion in the radial direction of biofilm growth; **(C)** The Y component is the range of deviation from biofilm-radial motion of landmarks. ***p = 0.0008, ****p < 0.0001; **(D)** A Geometric Density Histogram Ridgeplot of distance of movement (x-axis, “step-size”) between subsequent time-points of marked landmarks in time-lapse images of the indicated strains. The y-axis of each plot shows the frequency of occurrence of step sizes. Marked by black brackets are distinct populations of step-sizes in the NCIB 3610 (WT) images indicated by shoulders in the histogram. Strains used are WT NCIB 3610/NRS1473, *tapA* NRS3936/NRS6723, *sipW* NRS5488/NRS6718, *tasA* NRS5267/NRS6724, *tapA tasA* NRS5748/NRS6727, *epsH* NRS5906/NRS6728, *bslA* NRS2097/NRS5131.

Analysis of the rate of movement for the features tracked revealed further insight to the collective biomass motion (Fig. 5D, Fig. Movie S3 ‘s’). Calculating all the distances moved between each time-point, for NCIB 3610 three distinct populations of step-sizes are evident as shoulders in the density histogram (Fig. 5D, WT, black brackets), implying that there are different categories of motion. There was a similar pattern of shoulders in the distribution of *tapA* mutant strain step sizes as NCIB 3610, albeit with an increase in density of smaller step sizes (Fig. 5D *tapA*). The *sipW* mutant had a broad distribution of step-size with some features moving as far as NCIB 3610 with step-sizes up to almost 60 µm per 10 min interval but lost the three categories of step-sizes evident from NCIB 3610 and the *tapA* mutant strain (Fig. 5D *sipW*). The remainder of the matrix mutant strains displayed a unique profile of step-size distribution, generally showing much smaller step sizes than NCIB 3610 or the *tapA* and *sipW* mutant strains (Fig. 5D).

Finally, we tracked each landmark relative to the outer leading edge of its colony biofilm over time to determine if step-size correlated with the expansion rate of the colony biofilms (Fig. S5, Fig. Movie S3 ‘d’). Across each of the mutant strains, individual features either maintained pace with the expansion rate (approx. horizontal lines), travelled more slowly, increasing the distance from the expanding edge (positive gradient lines) or had an increased pace, catching up with the edge (negative gradient lines). Re-plotting these data to represent the change in distance to the edge over time, rather than the absolute distance, highlights each strain’s propensity to the movement categories described here (Fig. 6A). The predominant characteristic of NCIB 3610 feature trajectories is to fall behind the leading edge, as evidenced by the very few negative gradients when plotted (Fig. 6A). The trends of the *sipW* and *bslA* strains could be described similarly, but those of *tapA*, *tasA* and *epsH* strains include a higher proportion of negative gradients (Fig 6A). The variability in the trajectory travel profiles and landmark motion across the space implies that there are multiple forces acting on the biofilm biomass, and that during growth these forces push or pull the biofilm biomass in different directions in a matrix-dependent manner, revealing the impact of the matrix components on the material properties of the colony biofilm as a collective biomass.

**Figure 6.**
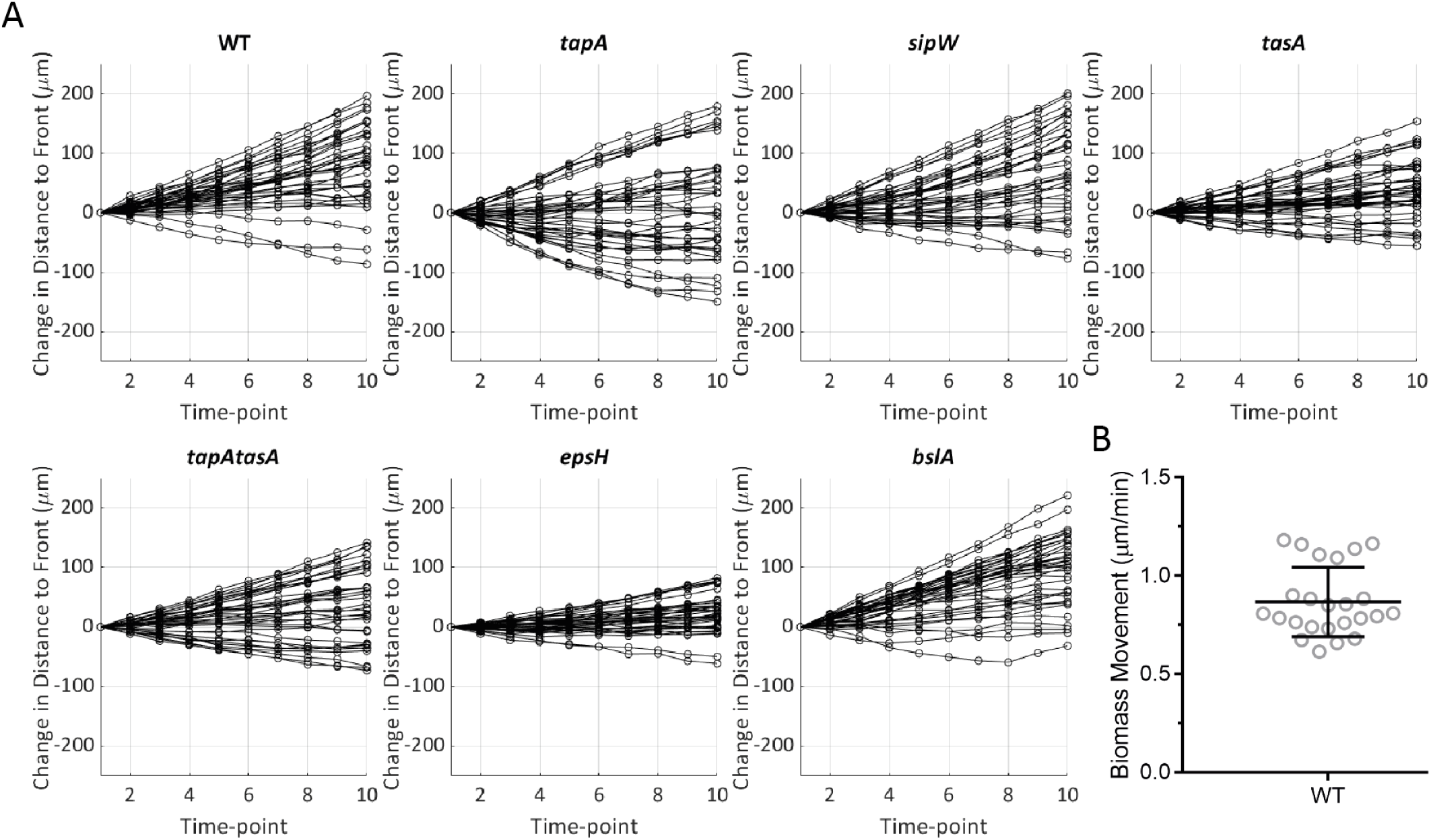
Detailing the dynamics of landmarks relative to the expanding edge of the biofilm. **(A)** Normalised graphs of Fig. S5 to the initial distance to the edge at t = 1. These show the change in distance since the first time point, highlighting the spread of landmark movement characteristics for each strain indicated (keeping pace with the edge (0), falling behind (+) or catching up (-) with the edge. Strains used are WT NCIB 3610/NRS1473, *tapA* NRS3936/NRS6723, *sipW* NRS5488/NRS6718, *tasA* NRS5267/NRS6724, *tapA tasA* NRS5748/NRS6727, *epsH* NRS5906/NRS6728, *bslA* NRS2097/NRS5131; **(B)** Rate of biomass flow at the biofilm-air interface. Visibly recognisable features (6 from each of 4 images across 2 biological repeats) of NCIB 3610 biofilms entering the image space were tracked by manual ROI placement for a duration of 15 hours and their mean rate of movement calculated.

### Colony biofilm expansion is a result of peripheral mono-layer growth

To address whether the collective movement of biofilm mass, from the interior of the community to the exterior, contributes to the expansive capability of *B. subtilis* colony biofilms or establish if expansion is driven by the outer edge of the community, we performed confocal imaging at regions immediately adjacent and interior to the areas of mono-layer chain expansion (Fig. S1C and D). We hypothesised that if movement of biofilm from the interior of the community contributed to expansion of the biofilm footprint, motion of the material moving towards the exterior would be at a rate that exceeds the rate of expansion directly measured for the biofilm periphery (Fig. 3C). Image collection started after 12 hours of biofilm growth and was sustained for a period of 15 hours. Data were acquired in two separate experimental set-ups capturing different perspectives of the colony biofilm growth: the biofilm-agar interface (Fig. Movie 5 WT; Fig. Movie S4) and biofilm-air interface (Fig. Movie 6 WT; Fig. Movie S5 A - F).

Imaging of the biofilm-agar interface from below, showed that NCIB 3610 and the *bslA* strain formed channels that hosted clusters of motile planktonic cells close to the expanding edge of the biofilm (Fig. Movie 5, WT and *bslA*; Fig. S1D). Both NCIB 3610 and the *bslA* mutant also had sectors of material in the growing colony biofilm material that showed concerted motion, first moving slowly in the outward direction before reversing towards the interior. The reverse motion is evident in biomass on the interior side of the field of view first, before appearing to stretch and drag the rest of the material towards the interior (Fig. Movie 5, WT and *bslA*). The NCIB 3610 colony biofilm material can be seen to press into the agar substratum, while the *bslA* biomass folds in on itself. This concerted motion within the biomass was not evident in the material formed by any of the other matrix deletion strains (Fig. Movie 5, compare NCIB 3610 and *bslA* to *tasA*) (Fig. Movie S4 A - E *tapA, sipW, tasA, tapAtasA, epsH*). Given that the indentations under biofilm peripheries were only observed in the cases of NCIB 3610 and the *bslA* mutant strain (Fig. 1C), it seems likely that, as movement at the agar interface was only observed in only these two strains, it is this movement that leads to formation of the indentations. Overall, no gross outward expansion of the biomass was apparent for any of the strains imaged at the agar interface (Fig. Movie 5, Fig. Movie S4 A - F).

Imaging of the biofilm-air interface (Fig. S1C) of NCIB 3610 showed transition of the peripheral mono-layer chains into a thicker tissue before the interior biomass entered and traversed the imaging field, moving towards the periphery of the colony biofilm at a rate of 0.87 µm/minute (Fig. 6B), (left to right, Fig. Movie 6, WT). In contrast, although each of the matrix deletion strains showed a transition of the peripheral mono-layer chains into a thicker tissue, the material displayed no coordinated translational motion for the 15-hour duration of imaging (Fig. Movie S5 A - F). When the mean outward movement of NCIB 3610 biomass at 0.87 µm/minute distant from the edge is compared to the mean peripheral expansion rate of 4.25 µm/minute (Fig. 3C) and is combined with the lack of any outward movement of the same biomass in the matrix deletion strains, we conclude that expansion of *B. subtilis* colony biofilms is primarily the consequence of growth at the biomass periphery.

### Thickening of mature biofilms requires the full suite of matrix components

Our final imaging dataset provided a more detailed perspective of the gross colony biofilm architecture by the application of confocal imaging to both the outer edge and the interior of NCIB 3610 colony biofilms. Both the fine structures seen in the periphery of the colony biofilm and the cylindrical structures of the colony biofilm interior at 48 hours of growth appeared to become thicker and more pronounced when equivalent regions were imaged at 72 hours of growth (Fig. 7A). To quantify this visual change, the images from the periphery were analysed for the total volume of material in the visible biomass, using the position of the agar as a reference (Fig. 7B). The analysis showed continued development of complex structures, alongside an ∼1.76-fold thickening of the biomass between 48-72 hours (Fig. 7C). In sharp contrast, similar imaging of the biofilm matrix mutants revealed few complex structures and no increase in biomass volume over a comparable time frame, mostly reporting a volume ratio very close to 1 (Fig. 7C, dashed line). The *bslA* mutant strain showed a slight reduction in volume, with a mean ratio of 0.87. Therefore, only wild-type NCIB 3610 colony biofilms have the propensity to increase in thickness (Fig. 7B, 7C), revealing the dependency on the full suite of biofilm matrix molecules for vertical expansion of the colony biofilm.

**Figure 7.**
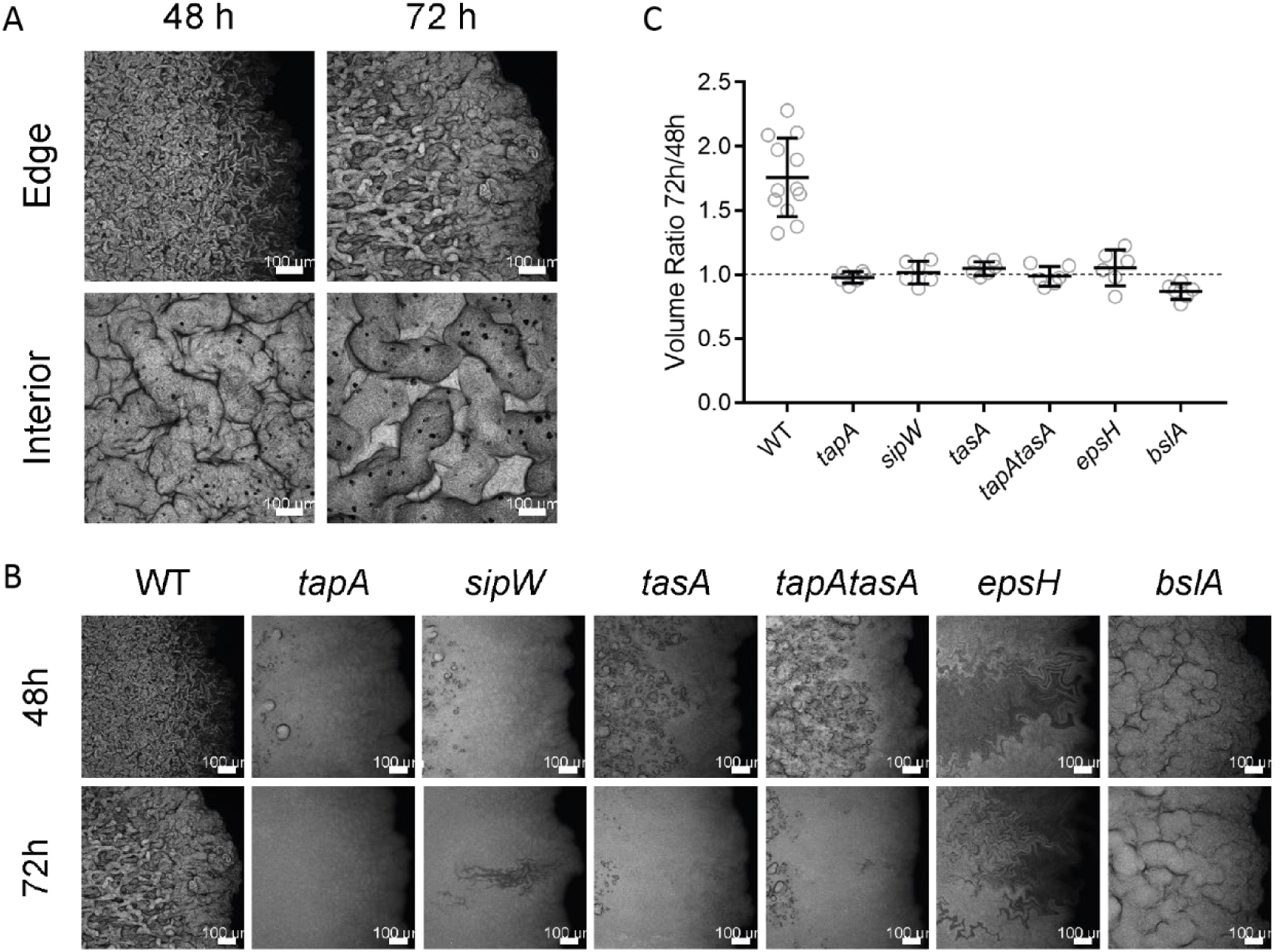
Maturation of biofilms. **(A)** Confocal imaging of wild-type NCIB 3610 biofilms expressing GFP at 48 hours and 72 hours after inoculation, grown at 30°C. Fields represent typical edge (top row) or interior (bottom row) morphologies becoming thicker and more pronounced over this period of growth. Maximum intensity projections are shown, scale bar is 100 µm; **(B)** GFP-Expressing Biofilms of wild-type and matrix mutants strains indicated were grown for 48 hours at 30°C before 3D images of their peripheries were acquired using CLSM (top row). The same biofilms were then grown for a further 24 hours and imaged again in similar locations (bottom row); **(C)** Volumetric images of biofilm peripheries of the strains indicated had their upper boundaries defined by segmentation in each xy location and the volume of the biomass calculated at 48 hours and 72 hours of growth at 30°C. Strains used are WT NRS1473, *tapA* NRS6723, *sipW* NRS6718, *tasA* NRS6724, *tapA tasA* NRS6727, *epsH* NRS6728, *bslA* NRS5131.

## Discussion

By developing and implementing time-lapse imaging and quantitation methods we observe and describe the dynamics of *Bacillus subtilis* colony biofilm formation, revealing the contribution of the extracellular matrix molecules. Through our analyses, we provide new insight to the microscale processes underpinning the formation of the resulting macroscale community. Understanding the specific function and impact of each of the polymeric matrix molecules secreted by cells in a colony biofilm is a challenging goal. By deleting the genes needed to produce each matrix component in turn (*tapA*, *sipW*, *tasA*, *epsH*, *bslA*), and in combination (*tapA tasA*), it was hoped that these functions would present themselves in the physical dynamics of the biomass. Our data show that BslA is not required to produce internal architecture leading to channel formation, but this feature is lost in the absence of *epsH* and any of the *tapA* operon gene products. Moreover, all the matrix component deletion strains showed a loss of coherence in the movement that accompanies growth.

### Matrix influence on cell division and physiology

Acknowledging that cells undergoing sessile growth experience physiological conditions that are different from those in planktonic culture (Dar et al., 2021), we developed a new method to calculate the cell division time of bacteria growing in the context of a biofilm with single-cell resolution. Each of the matrix-deficient strains, except the *epsH* mutant, divide at comparable rates to NCIB 3610 during expansion of cell chains at the periphery of the colony biofilm. That the absence of exopolysaccharide results in increased cell-cycle times helps to explain the *eps* mutant colony biofilms’ diminutive form, but the mechanism of this deficiency is unexplained and warrants further investigation.

There is a disparity in observed doubling times at different stages and physical locations in biofilm growth. During initial colonisation of the agar, in the period after deposition of the cells, NCIB 3610 had a mean doubling time of around 4.3 hours (Fig. 2C), while after 12 hours of growth the chains of cells at the expanding periphery double every 54 minutes, on average (Fig. 4B). We speculate that the physiology of the cells is very different in these two cases: during colonisation the cells are switching from planktonic to sessile growth mode and are ultimately constrained within the area of inoculation; at the periphery at later times the cells have adapted to sessile growth and have immediate access to free space in which to grow. Given the range of morphological structures in NCIB 3610 colony biofilms, from dense regions of cells, aerial projections, and fluid-filled cavities, we surmise that there are many physiological pressures present that will lead to regulatory adaptations within a local area of cells.

Altered gene regulation in micro-environments of the biofilm may also account for the bimodality of expansion rates measured for the *sipW* strain, where two distinct clusters of speeds were detected in the fields of view that were selected for imaging (Fig. 3C). SipW is a signal peptidase responsible for processing and releasing nascent TapA and TasA from the cell (Tjalsma et al., 2000), but also has a matrix gene regulatory function in the context of biofilms (Terra et al., 2012). It is conceivable that disturbing the balance of matrix gene regulatory signals in situations of heterologous physiological pressures could result in inconsistent responses of cells in a local area. However, on a whole-biofilm scale, *sipW* gene deletion leads to reduced morphological complexity and smaller biomass footprints (Fig. 1A and B) compared to NCIB 3610.

### Matrix influence on tissue motion and structure

Described as an active matter system, chains of *B. subtilis* cells on an agar substrate have been observed to behave nematically and produce topological defects to alleviate stresses incurred by growth, creating 3-dimensional structures (Yaman et al., 2019). When considering why mutant strains deficient in matrix components form biofilms with smaller footprints and display less complex morphology, we find that it is not the initial growth rate of the cells, as they transition into sessile growth from planktonic culture, that is the root cause of macroscale impact. Rather, we observe 2D chains growing into 3D structures a short distance from the edge of expanding biofilms, and we propose that the dynamism of movements of these ordered structures away from the purely lateral direction is dependent on strength provided by the combination of matrix molecules present. This tensile stress is transmitted over a larger length-scale when the full suite of matrix molecules is produced, confirmed by the deviations of feature motions from the radial direction of biofilm growth, and the formation of structures that extend both aerially from the biomass and embed into the agar substratum. This process produces and maintains internal cavities hosting fluid and pockets of swimming planktonic cells that contribute to heterogeneity in the isogenic community (Vlamakis et al., 2008). The matrix-mutant biofilms show a lack of 3-dimensional organised structures. If this is due to a loss of scaffolding in the spacing between chains of cells, it follows that there might be tighter cell packing, that the overall surface area is reduced and total cell density greater in these cases. Tighter cell packing in this way has negative implications with regards to nutrient and oxygen permeation (Warren et al., 2019), and would go some way to explaining why matrix mutant biofilms with cell division rates similar to NCIB 3610 at the periphery produce consistently smaller biofilms.

### Mode of biofilm growth

Osmotic spreading, or the swelling of the *B. subtilis* colony biofilm by uptake of water from the substratum, has been proposed (Seminara et al., 2012). However, by directly observing the motion of biomass at various locations throughout biofilm expansion we see that instead of swelling from within, NCIB 3610 colony biofilms have the highest rate of lateral growth at the periphery, and this growth results from cell division in mono-layer chains rather than biomass accumulating and pushing from the interior. Moreover, for NCIB 3610 the slow biomass movement inside the biofilm towards the exterior was only observed at the biofilm-air interface and not at the biofilm-agar interface. This model of colony biofilm expansion is consistent with waves of biofilm matrix gene expression at the periphery of the biofilm in *B. subtilis* (Srinivasan et al., 2018) and zones of active growth in similar colony biofilm communities formed by *Escherichia coli* (Rooney et al., 2020). Furthermore, motion of material at the underside (agar proximal) of the colony biofilm, that was initially at a short distance from the edge, indicates that as growth continues this foundational biomass not only lacks expansive sliding, but also becomes embedded into the agar substratum, forming peripheral indentations in the substrate (Zhang et al., 2015).

### Overarching Conclusion

*In toto,* this study has provided unprecedented views of a complex macroscale microbial community forming from an initial dispersed population of cells. The synergy of the matrix molecules in providing the tensile strength needed to form the complex architecture becomes apparent, and collective motion of the community reveals how the forces generated by cell growth are dissipated in the material. Whilst the specific functions of each matrix component remain elusive, in total, the matrix components function with dependence on each other to confer structure and strength, as indicated by the mutant strains being unable to maintain even small-scale morphological features or to increase in thickness as the biofilm matures. Combining imaging analysis of this nature with microscale selection of distinct micro-pockets of cells would allow the molecular processes underpinning the transition from the motile state to the embedded social community to be followed. In the future it will be intriguing to apply these combined techniques to identify the benefits of matrix molecules on a broad range of bacterial species in their varied natural environments.

## Acknowledgements

Work in the NSW, FAD and CEM laboratories is funded by the Biotechnology and Biological Science Research Council (BBSRC) [BB/P001335/1, BB/R012415/1]. We are grateful to Dr David J. Williams for visualisation of data in R, Dr Ryan Morris, and Dr Lukas Eigentler for thoughtful comments on the manuscript, and other members of the Stanley-Wall lab for helpful discussions. We would like to acknowledge the Dundee Imaging Facility at the University of Dundee and note that part of the work presented here has been published in the doctoral thesis of Michael Porter.

## Data availability statement

Computational code is available on Github and has been archived by Zenodo - 10.5281/zenodo.6563834.

The experimental datasets have been archived using BioStudies (Sarkans et al., 2018) with accession S-BIAD474 and can be found at https://www.ebi.ac.uk/biostudies/studies/S-BIAD474

## Author Contributions using CRediT

Conceptualisation: MP, NSW, CEM

Data curation: MP

Formal Analysis: MP

Funding acquisition: NSW, CEM, FAD

Investigation: MP

Methodology: MP

Project administration: MP

Resources: MP

Software: MP

Supervision: NSW

Validation: MP

Visualization: MP

Writing – original draft: MP, NSW

Writing – review & editing: MP, NSW, CEM, FAD

## Materials and Methods

### Construction of Strains

Strains generated for this study that carried genes encoding GFPMut2 or mKate2 under constitutive promoters were produced by SSP1 bacteriophage transduction of genetic material from previously published strains (see Strain Table). Briefly, bacteriophages were generated from the donor strain carrying the desired genetic element (*sacA*::P*spachy-gfp-kan* or *amyE*::P*spachy-mKate2-spc*) using previously published methods and introduced into the recipient strain (Verhamme et al., 2007).

### Construction of FtsZ-GFP Fusion Protein Strains

A marker-less *ftsZ-gfp* fusion gene was created at the native *ftsZ* locus by homologous recombination using vector pMAD2 (Arnaud et al., 2004) containing the gene for GFPMut2 flanked by homology arms (HA) of *ftsZ* at the 5’ end and *bpr* at the 3’ end. The two HAs were created by PCR of NCIB 3610 genomic DNA using primers NSW3027 and NSW3033, and NSW3037 and NSW3032 respectively (see Primer Table). A *gfpmut2* fragment was generated by PCR from genomic DNA of strain NRS2388 (see Strain Table) using primers NSW3034 and NSW3035. These PCR fragments contained flanking homology for assembly using a Gibson Assembly kit (New England Biolabs) to yield the final vector pMAD2-5’HA-*gfpmut2*-3’HA (pNW2410) which was verified by sequencing using primers NSW3023 and NSW3024. This vector was introduced into *B. subtilis* strain 168 cells, then into final recipient strains (NCIB 3610, NRS3936, NRS5488, NRS5267, NRS5748, NRS5906 and NRS2097) (see Strain Table) via phage transduction (Verhamme 2007). Integration of *ftsZ-gfpmut2* into the genomes of these strains was achieved by a method published previously (Arnaud et al., 2004) and positive colonies were screened for GFP fluorescence on a stereomicroscope and PCR analysis. The resultant strains had the addition of constitutively expressing mKate2 at the *amyE* locus via phage transduction (Verhamme et al., 2007) from strain NRS5841, selecting on spectinomycin (100 µg/ml) plates to yield final strains NRS6781, NRS6793, NRS6794, NRS6795, NRS7525, NRS7526 and NRS7527.

### Culture of Cells and Biofilm Growth

Colony biofilms were founded from a pre-culture grown in 3 ml liquid Lysogeny Broth (LB, per litre: 10 g yeast extract, 5 g NaCl, 10 g tryptone) from a single colony streaked from frozen strain stocks on an LB agar (1.5% w/v SelectAgar, Invitrogen) plate. The liquid cultures were incubated for three to four hours at 37°C shaking at 200 rpm and adjusted to the same optical density (OD_600_) as the lowest culture in a set, in the range of OD_600_ of 0.9 to 1. Strains expressing fluorescent molecules were then mixed with their parental strains (see Strain Table) after normalisation of cell density (80 µl non-fluorescent strain with 20 µl fluorescent strain). Colony biofilms were inoculated from this mixture by spotting 1 µl of the pre-culture onto MSgg (5 mM potassium phosphate and 100 mM MOPS at pH 7.0 autoclaved and cooled to 55°C then supplemented with 2 mM MgCl_2_, 700 μM CaCl_2_, 50 μM MnCl_2_, 50 μM FeCl_3_, 1 μM NaCl_2_, 2 μM thiamine, 0.5 % (v/v) glycerol, 0.5 % (v/v) glutamic acid) agar (1.5% w/v). The pre-culture spot was air dried under a flame for 5 minutes before incubating the covered colony biofilms at 30°C in a humidified box or transferring them immediately to a pre-warmed microscope chamber.

### Colony biofilm and indentation imaging

Macro imaging of whole biofilms and areas of agar indentation were performed on a Leica stereoscope (M205FCA) using reflected light from a ring illuminator and either a 0.5x 0.2 NA or 1x 0.2 NA objective.

### Confocal imaging of constitutive GFP colony biofilms

LabTek II 2-chamber microscope vessels (Thermo Scientific) were used to grow and image biofilms 30 minutes, 10.5 hours, 48 hours, and 72 hours after cell deposition (as indicated), with 2 biofilms per chamber. Two vessels could be imaged concurrently, totalling 8 biofilms. For imaging the biofilm-agar interface, the chambers were filled with 750 µl MSgg agar to put the biofilm into the working distance of the objective without excessively scattering light with the agar, in all other cases 4.4 ml MSgg agar was used, leaving a small headspace for the biofilms to grow, and be imaged from the biofilm-air interface. After solidifying, the MSgg agar was air-dried in a laminar flow hood for 45 minutes.

Non-fluorescent and constitutive GFP-expressing equivalent strains of wild-type and matrix gene mutants (see Strain Table) were grown, normalised for optical density and mixed, as detailed above. The cells were placed onto 4 ml MSgg agar (1.5% w/v) set into each chamber of LabTek II 2-chamber microscope vessels (Thermo Scientific) set into various containers, as detailed in specific methods. Two vessels were prepared per experiment, giving four chambers, and the MSgg agar was dried in a flow hood for 45 minutes before spotting. Two biofilms per strain analysed were spotted in a single chamber, spaced evenly apart.

A Leica SP8 upright confocal was used with a chamber pre-warmed to 30°C. The 488 nm argon laser was set to 2% power and rapid imaging was performed with a resonant mirror at 8 kHz with bi-directional scanning and line averaging of 16x. Gain of the PMT detector was set to between 550 v and 650 v to avoid saturating pixel values. The pinhole was set to 1 AU for a wavelength of 525 nm. Nyquist acquisition was achieved using a 10x, 0.3 NA long working distance objective for z-stacks with 3.87 µm steps and a scan of 1024 x 1024 pixels over an 890 µm area. Time-lapse imaging was set with an interval of 10 minutes. For imaging the biofilm-agar interface the vessels had their lids removed and were inverted onto the microscope stage. For imaging the biofilm-air interface the vessels were placed on the stage and had their lids replaced with a 60 mm x 24 mm #1.5 coverglass to prevent biofilm exposure to the flowing heated air of the microscope chamber. The biomass proximal to the petri dish edge was imaged.

### Confocal imaging of FtsZ-GFP fusion strains

Vessels were prepared by adhering a single Gene Frame (65 µl) (Thermo Scientific) onto a glass microscope slide, leaving the plastic aperture cover in place. A stepped flat mould was created by gluing 22 mm x 22 mm glass coverslips to a second microscope slide at a distance apart that would align to the inside edges of the Gene Frame. 150 µl of MSgg agar (at 55°C) was pipetted into the aperture of the Gene Frame, avoiding forming bubbles on the surface, and the stepped mould was immediately inverted over the aperture and held in place for 30 s to allow the agar to set, slightly raised above the height of the Gene Frame, and removed by gently sliding off. This was done to ensure that the sample could still be mounted after agar shrinking during incubation before imaging. After mixing cultures (see above), three biofilms were inoculated onto the agar, spaced equally. The slide was incubated in a humidified box at 30 °C for 12 hours before mounting. To maximise oxygen availability during imaging, most of the agar in the Gene Frame was cut away with a sterile scalpel, leaving approximately one quarter of each biofilm (Fig. S2Biii) before peeling off the aperture cover from the Gene Frame and applying half a gas-permeable polymer coverslip (Ibidi) that had been cut in two equal pieces.

Imaging was performed on a Leica SP8 inverted confocal microscope with a chamber pre-warmed to 30 °C. Excitation of GFP and mKate2 was achieved with 488 nm and 568 nm lasers, respectively, set to 2% and 0.1% power and read by hybrid detectors with a gain of 10 v and 2 v. The pinhole was set to 1 AU at 525 nm for optimal resolution of GFP and both channels were acquired simultaneously. Fast imaging was employed with a resonant mirror at 8 kHz and line averaging of 16x and the image field was 752 x 752 pixels. Three image fields were set around the periphery of each of the three biofilms on the slide, generating nine movies per acquisition, with a z-stack specified to capture at Nyquist sampling with the 63x 1.4 NA objective. Images were collected every 10 minutes for 5 hours.

### Image handling and figure preparation

All microscopy data were imported into an OMERO server (Allan et al., 2012) for organisation, annotation, and analysis. Maximum intensity image projection and annotation with ROIs were performed with OMERO.insight. Static image figures were created using OMERO.figure while movie figures were prepared using bespoke code in Matlab (MathWorks) (See Image Analysis). Raw, projected and ROI-annotated image data are available at https://www.ebi.ac.uk/biostudies/studies/S-BIAD474 which contains a link to live data on the OMERO server.

### Image analysis

Analysis of intensity images and ROI positions were performed in Matlab (R2019a) using the OMERO.matlab toolkit (v5.5.4) to read data from the OMERO server (v5.6.3). All code is available at (doi: 10.5281/zenodo.6563834). A summary of analysis workflows is as follows:

#### Initial doubling time

three identical square ROIs were placed on images in the internal region of the newly-spotted biofilm and propagated through every time-point of the images. The ROIs were used to define patches of the image to be downloaded, and the mean intensity of each patch was calculated. The means were normalised to the highest value over all time-points, producing a sigmoid curve. The highest gradient was detected and used to define the doubling time.

#### Expansion rate

Time-lapse images of chains of cells, oriented to move left to right across the image space, were segmented using Otsu thresholding (Otsu, 1979) until the chains reached the far edge of the image. The right-most pixel segmented had its x-axis location recorded for each time point, converted from pixel units to µm using the pixel size metadata. These position data were plotted, and a straight line was fitted with high confidence. Expansion rate was calculated from the gradient of the fitted line.

#### Tracking of feature landmarks

For each time-lapse image, six visibly recognisable features were marked with a point-ROI, which was propagated to the time-point where the chains reached the far edge of the image. Features were chosen to be spread across the height and width of the observed chains in the image at the first time-point. The locations of the points were then updated manually for each time-point. For fair comparison, the number of time-points used for downstream analysis was limited to that of the shortest tracks detected. The XY location of each point was used to generate tracks across the image space, from which the step-sizes (distance moved between time-points), distance to the edge (using the edge location from Expansion rate, above) and speed were calculated.

#### FtsZ cycle time

Time-lapse imaging was examined for the appearance of new Z-rings. Upon first sighting, a Z-ring was marked with a single point ROI and designated as the ‘root’ of a pedigree with a ‘0’ entered as the ROI’s comment. Progeny were followed over time and when new daughter Z-rings became visible they were marked with a point and given the id of their parent’s ROI as a comment. The pedigrees were followed over multiple generations, using the constitutive mKate2 signal of this strain to visually delineate daughter cells. Each image’s ROIs were then analysed in Matlab, sorted into pedigrees using the root (‘0’) and parent ROI id comments to form links between them, calculating the interval from the difference in time-point of the ROIs.

#### Footprint size analysis

The area occupied by whole biofilms was calculated in Matlab by segmentation using Otsu thresholding (Otsu, 1979) of reflected-light images acquired on a stereomicroscope. Pixel units were converted to µm using pixel size metadata from the image and reported as µm^2^.

#### Biofilm thickness ratio

Volumetric images from the edge of biofilms were segmented using Otsu thresholding (Otsu, 1979), and for each XY coordinate the upper-most pixel in the Z dimension containing segmented data was recorded. The total volume of biomass was then calculated using the z-section size metadata and a ratio was generated between biomass volumes at 48 hours and 72 hours of growth.

#### Internal biomass movement

Visibly recognisable structures moving across the image field were identified and marked with a point ROI that was used to manually track progress across the images for a minimum duration of 11 hours. The Euclidean distance given by the change in coordinates of the point ROIs during each 18-minute time-interval was then used to calculate movement rate.

#### Movie figure preparation

Time-lapse images were downloaded into Matlab plane by plane from OMERO to generate frames onto which ROIs could be used to highlight specific features. Scale bar lengths were chosen and drawn onto each frame using the pixel size metadata of each image. The frames were then collated and saved as .avi files with a framerate of 2 per second.

### Graphing and Statistical Analysis

Histogram ridge plot of step-sizes was generated in R from data exported from Matlab, code for which is available at (doi: 10.5281/zenodo.6563834). Other graphs were drawn in Matlab and Prism (Graphpad). Statistical analyses were performed in Prism and p-values reported are the results of 1-way ANOVA using Dunnett’s multiple comparisons test with significance set to p = 0.05.

---

**Table 1:** Strains used in this study

| Strain | Relevant Genotype/Description † | Source*/Construction** |
| --- | --- | --- |
| 168 | *trpC2* | *Bacillus* Genetic Stock Centre |
| NCIB 3610 | Prototroph | *Bacillus* Genetic Stock Centre |
| NRS1113 | *amyE::Pspachy-gfpmut-gfp::cat::spc* | (Stanley et al., 2003) |
| NRS1471 | *trpC2 pheA1 sacA-Pspachy-gfp (kan)* | (Verhamme et al., 2007) |
| NRS1473 | 3610 *sacA-Pspachy-gfp (kan)* | (Hobley et al., 2013) |
| NRS2097 | 3610 *bslA::cat* | (Ostrowski et al., 2011) |
| NRS2388 | 168 *sacA::PtapA-gfp::kan* (published as *PyqxM-gfp*) | (Ostrowski et al., 2011) |
| NRS3936 | 3610 Δ*tapA* | (Erskine et al., 2018) |
| NRS5131 | 3610 *bslA::cat, amyE::Pspachy-gfpmut-gfp::cat::spec* | SPP1 NRS1113 → NRS5488 |
| NRS5267 | 3610 Δ*tasA* | (Erskine et al., 2018) |
| NRS5488 | 3610 Δ*sipW* | (Earl et al., 2020) |
| NRS5748 | 3610 Δ*tapA* Δ*tasA* | (Erskine et al., 2018) |
| NRS5841 | 3610 Δ*comI amyE::Phyperspank-mKate2 (spc)* | (van Gestel et al., 2014), re-stocked as NRS5841 |
| DS6776/NRS5906 | 3610 Δ*epsH* | (Roux et al., 2015), re-stocked as NRS5906 |
| NRS6718 | 3610 Δ*sipW, sacA::Pspachy-gfp (kan)* | SPP1 NRS1471 → NRS5488 |
| NRS6723 | 3610 Δ*tapA, sacA::Pspachy-gfp (kan)* | SPP1 NRS1471 → NRS3936 |
| NRS6724 | 3610 Δ*tasA, sacA::Pspachy-gfp (kan)* | SPP1 NRS1471 → NRS5267 |
| NRS6727 | 3610 Δ*tapA, \Delta*tasA, sacA::Pspachy-gfp (kan)** | SPP1 NRS1471 → NRS5748 |
| NRS6728 | 3610 Δ*epsH, sacA::Pspachy-gfp (kan)* | SPP1 NRS1471 → NRS5906 |
| NRS6778 | 3610 *ftsZ-gfp* | SPP1 NRS6786 → NCIB 3610 |
| NRS6781 | 3610 *ftsZ-gfp, amyE::Phyperspank-mKate2 (spc)* | SPP1 NRS5841 → NRR6778 |
| NRS6786 | 168 *pMad2-ftsZ-gfp* | *pMad2-ftsZ-gfp* → 168 |
| NRS6787 | 3610 Δ*sipW, ftsZ-gfp* | SPP1 NRS6786 → NRS5488 |
| NRS6788 | 3610 Δ*tasA, ftsZ-gfp* | SPP1 NRS6786 → NRS5267 |
| NRS6789 | 3610 Δ*tapA \Delta*tasA, ftsZ-gfp** | SPP1 NRS6786 → NRS5748 |
| NRS6790 | 3610 Δ*epsH, ftsZ-gfp* | SPP1 NRS6786 → NRS5906 |
| NRS6791 | 3610 *bslA::cat, ftsZ-gfp* | SPP1 NRS6786 → NRS2097 |
| NRS6792 | 3610 Δ*tapA, ftsZ-gfp* | SPP1 NRS6786 → NRS3936 |
| NRS6793 | 3610 Δ*tapA, ftsZ-gfp, amyE::Phyperspank-mKate2 (spc)* | SPP1 NRS5841 → NRS6792 |
| NRS6794 | 3610 Δ*sipW, ftsZ-gfp, amyE::Phyperspank-mKate2 (spc)* | SPP1 NRS5841 → NRS6787 |
| NRS6795 | 3610 Δ*tasA, ftsZ-gfp, amyE::Phyperspank-mKate2 (spc)* | SPP1 NRS5841 → NRS6788 |
| NRS7525 | 3610 Δ*tapA, \Delta*tasA, ftsZ-gfp, amyE::Phyperspank-mKate2 (spc)** | SPP1 NRS5841 → NRS6789 |
| NRS7526 | 3610 Δ*epsH, ftsZ-gfp, amyE::Phyperspank-mKate2 (spc)* | SPP1 NRS5841 → NRS6790 |
| NRS7527 | 3610 *bslA::cat, ftsZ-gfp, amyE::Phyperspank-mKate2 (spc)* | SPP1 NRS5841 → NRS6791 |

‡Drug resistance cassettes are as follows: kan, kanamycin resistance; spc, spectinomycin resistance; cat, chloramphenicol resistance.
\*Strains sourced from the *Bacillus* Genetic Stock Centre or the indicated publication.
\*\* Strain construction is denoted as DNA or SPP1 bacteriophage from donor strain transformed or transduced into recipient strain following the direction of the arrow.

**Table 2:**
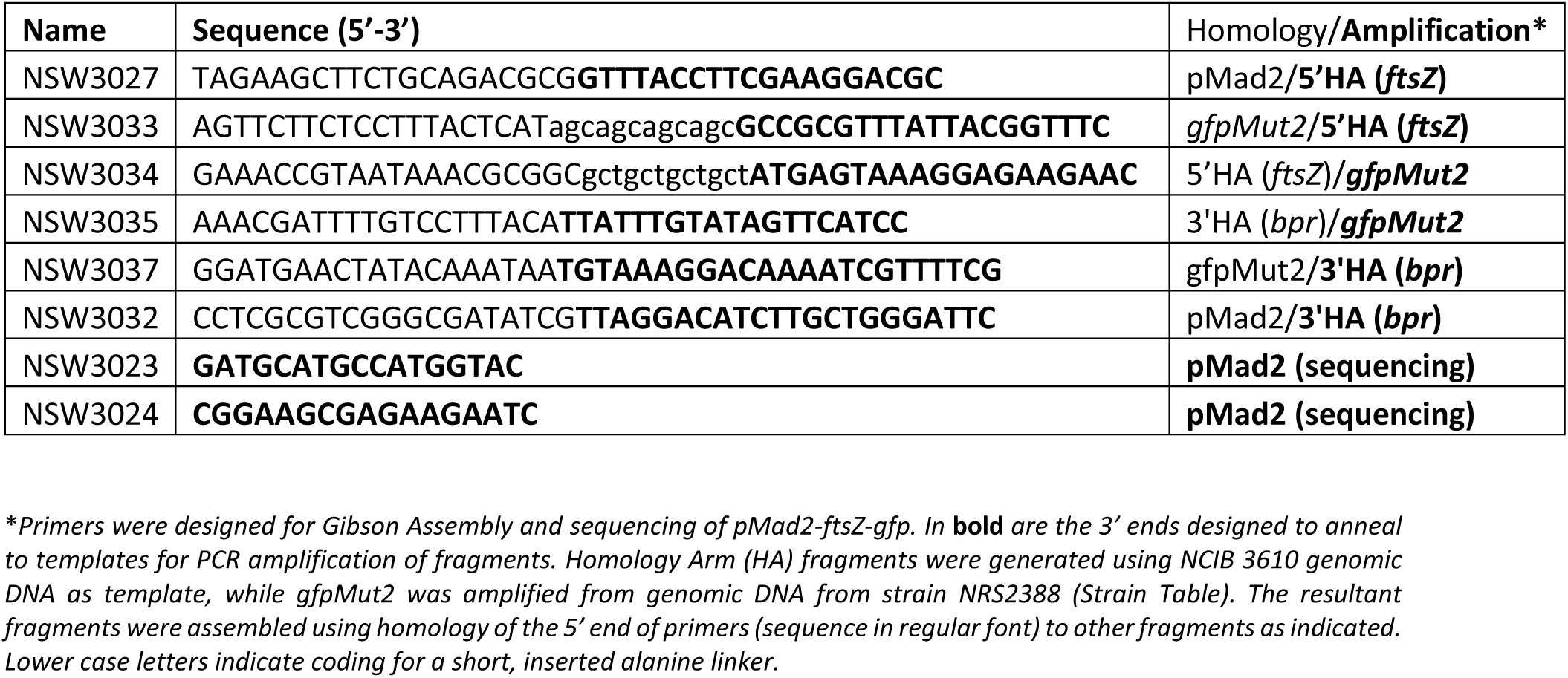
Primers used in this study

**Figure Movie 1.**
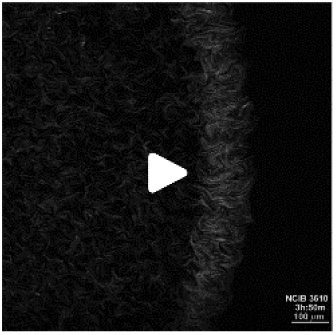
Initial cell growth. Time-lapse confocal microscopy of strains NCIB 3610 (80% population) and GFP-producing NCIB 3610 derivative NRS1473 (20% population) cocultured from 30 minutes after cell deposition. Z-stacks were acquired every 10 minutes for 10 hours, and a maximum intensity projection is presented. The timestamp is the approximate age of the biofilm since cells were deposited and the scale bar is 100 µm.

**Figure Movie 2.**
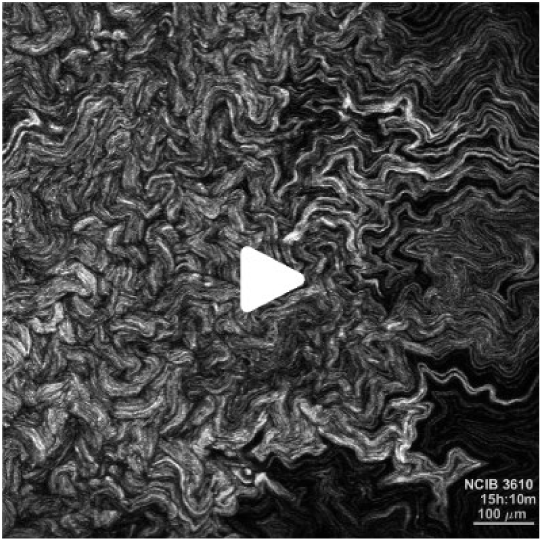
Cell chains emanate from the biofilm periphery. Time-lapse confocal microscopy of strains NCIB 3610 (80% population) and the NCIB 3610 GFP-producing derivative NRS1473 (20% population) from 10.5 hours after cell deposition and incubation at 30°C. Z-stacks were acquired every 10 minutes for 14 hours, and a maximum intensity projection is presented. The timestamp is the approximate age of the biofilm since cells were deposited and the scale bar is 100 µm.

**Figure Movie 3.**
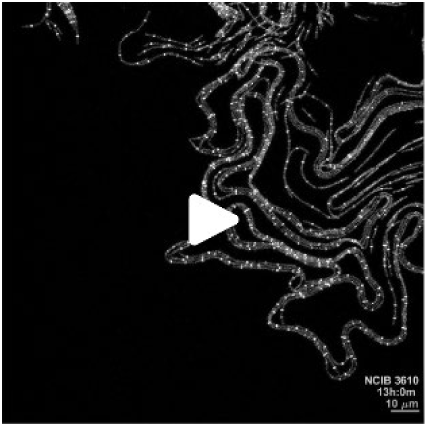
Example of single cell division time within the context of a colony biofilm. Time-lapse confocal microscopy of mixed strains NCIB 3610 (80%) and FtsZ-GFP-producing NRS6781 (20%) from 12 hours after cell deposition and incubation at 30°C. Z-stacks were acquired every 10 minutes, and a maximum intensity projection is presented. The timestamp is the approximate age of the biofilm since cells were deposited and the scale bar is 10 µm

**Figure Movie 4.**
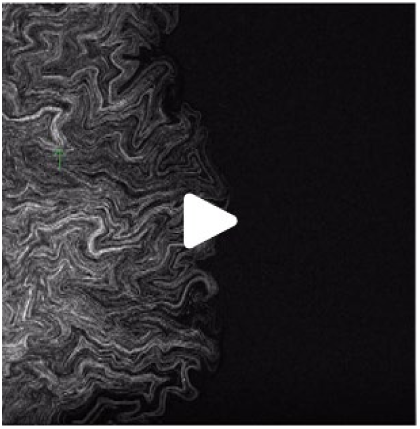
Example of biomass landmark motion. Time-lapse confocal microscopy of mixed strains NCIB 3610 (80%) and GFP-producing NRS1473 (20%) from 10.5 hours after cell deposition and incubation at 30°C. Z-stacks were acquired every 10 minutes for 14 hours, and a maximum intensity projection is presented. Green arrows highlight aerial structures that form a short distance from the expanding edge of the biofilm.

**Figure Movie 5.**
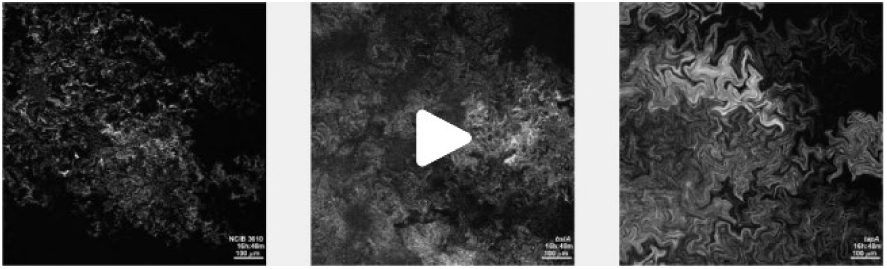
Biomass structure development at the biofilm-agar interface. Time-lapse confocal microscopy at the biofilm-agar interface of mixed strains of the genotypes stated containing 20% GFP-producing cells. Imaging was begun at 12 hours after cell deposition and incubation at 30°C. Z-stacks were acquired every 18 minutes, and a maximum intensity projection is presented. The timestamp is the approximate age of the biofilm since cells were deposited and the scale bar is 100 µm. Strains used are NCIB 3610/NRS1473, *bslA* NRS2097/NRS5131, *tasA* NRS5267/NRS6724.

**Figure Movie 6.**
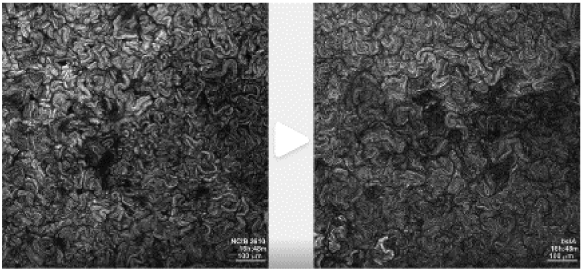
Biomass movement at the biofilm-air interface. Time-lapse confocal microscopy at the biofilm-air interface of mixed strains of the genotypes stated containing 20% GFP-producing cells. Imaging was begun at 12 hours after cell deposition and incubation at 30°C. Z-stacks were acquired every 18 minutes, and a maximum intensity projection is presented. The timestamp is the approximate age of the biofilm since cells were deposited and the scale bar is 100 µm. Strains used are NCIB 3610/NRS1473, *bslA* NRS2097/NRS5131

## Supplemental Figures

**Figure S 1.**
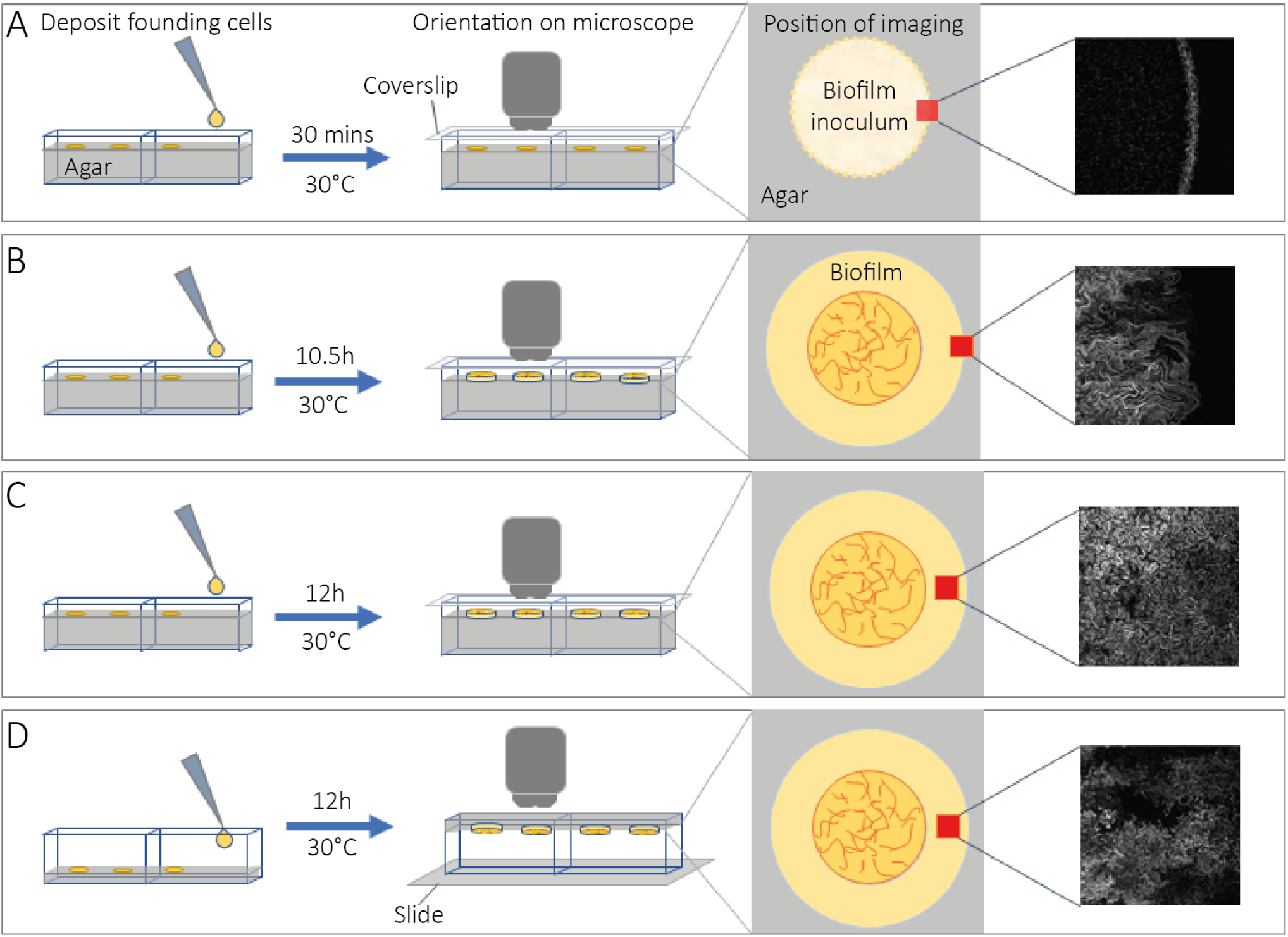
Schematic of imaging experiments. Time-lapse confocal microscopy was performed on biofilms from different perspectives and at various points in development. **(A - C)** Lab-Tek II chambers were filled with MSgg agar (see Methods) to leave a small headspace for the biofilm to grow; **(D)** Lab-Tek II chambers were covered with a thin pad of MSgg agar to allow visualisation of the underside of the biofilm. In all cases cells were deposited and the chambers were incubated before imaging for: 30 minutes (A); 10.5 hours (B); or 12 hours (C, D). Lab-Tek lids were removed and replaced with a long coverslip to protect the biofilm from warm airflow in the microscope chamber while allowing imaging of the biofilm-air interface (A – C) or inverted onto a microscope slide for imaging of the biofilm-agar interface (D). Fields of view were set to capture the edge of the biomass (A, B) or a region entirely within the biomass but near to the edge (C, D).

**Figure S 2.**
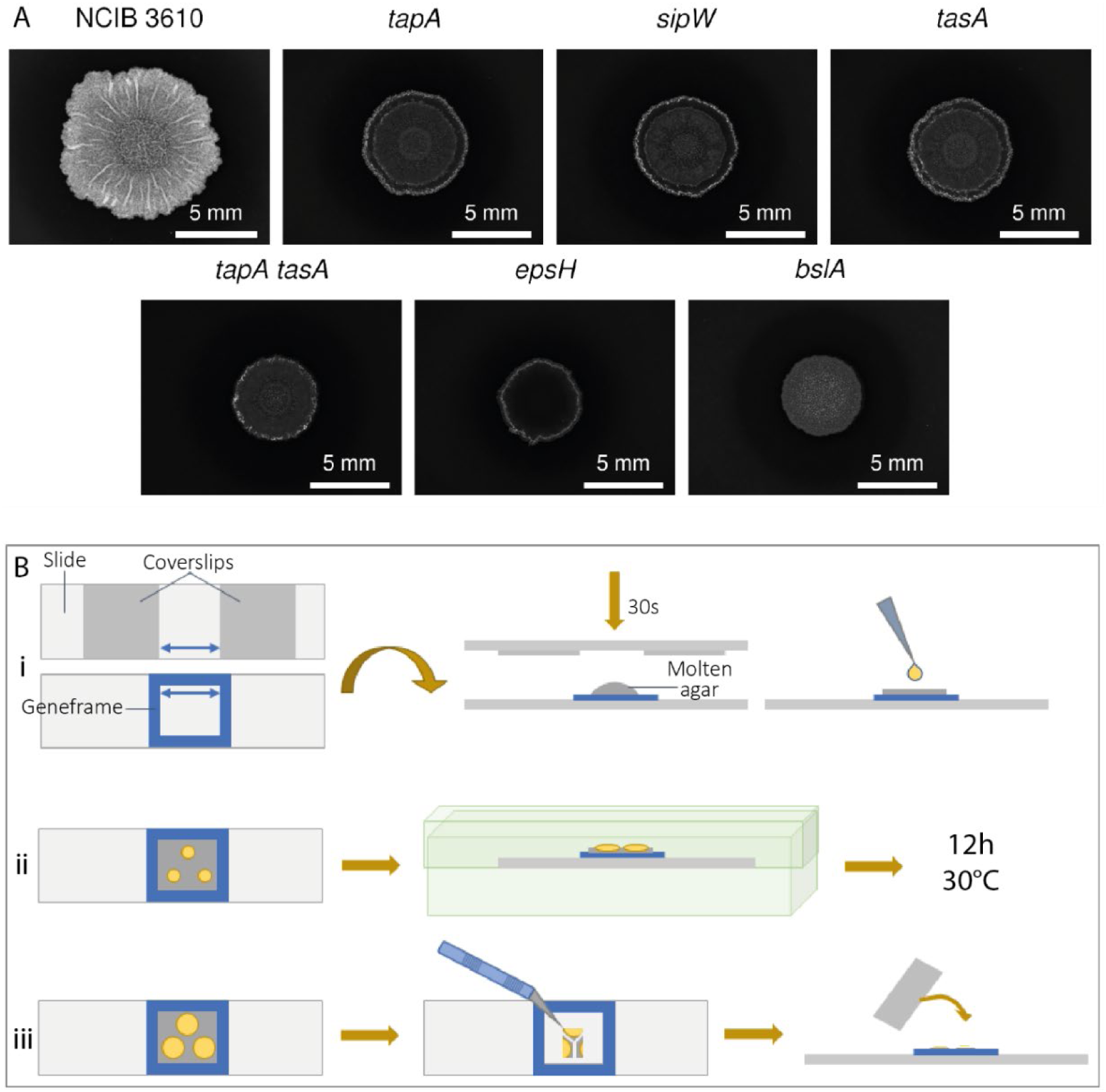
Experimental summary for strains expressing ftsZ-gfp. **(A)** Wild type and matrix mutant strains producing the FtsZ-GFP fusion protein and the fluorophore mKate2 display morphologies akin to their parent strains. Strains shown are: NCIB 3610 background NRS6781; *tapA* NRS6793; *sipW* NRS6794; *tasA* NRS6795; *tapA tasA* NRS7525; *epsH* NRS7526; *bslA* NRS7527. The colony biofilms were grown at 30°C for 48 hours prior to imaging; **(B)** Experimental setup for high resolution imaging of strains expressing *ftsZ-gfp*: (i) An adhesive Geneframe was attached to a glass side, and a flat mould was prepared by affixing two coverslips to a second slide, spaced apart to match the width of the aperture of the Geneframe (blue arrows). Molten MSgg agar was pipetted into the Geneframe and immediately flattened with the mould, held down for 30 seconds to set. Cells were deposited onto the agar pad, which is slightly raised relative to the edge of the Geneframe to allow for shrinkage during incubation. (ii) After inoculating three biofilms onto the agar, the slide was placed into a humidified box (microtube storage box with water in the outer positions) and incubated at 30°C for 12 hours. (iii) Immediately prior to imaging, the slide was removed from the box and most of the agar and unnecessary biomass was cut away with a scalpel to increase air space and reduce the demand for oxygen. A gas-permeable coverslip was applied to complete mounting of the sample.

**Figure S 3.**
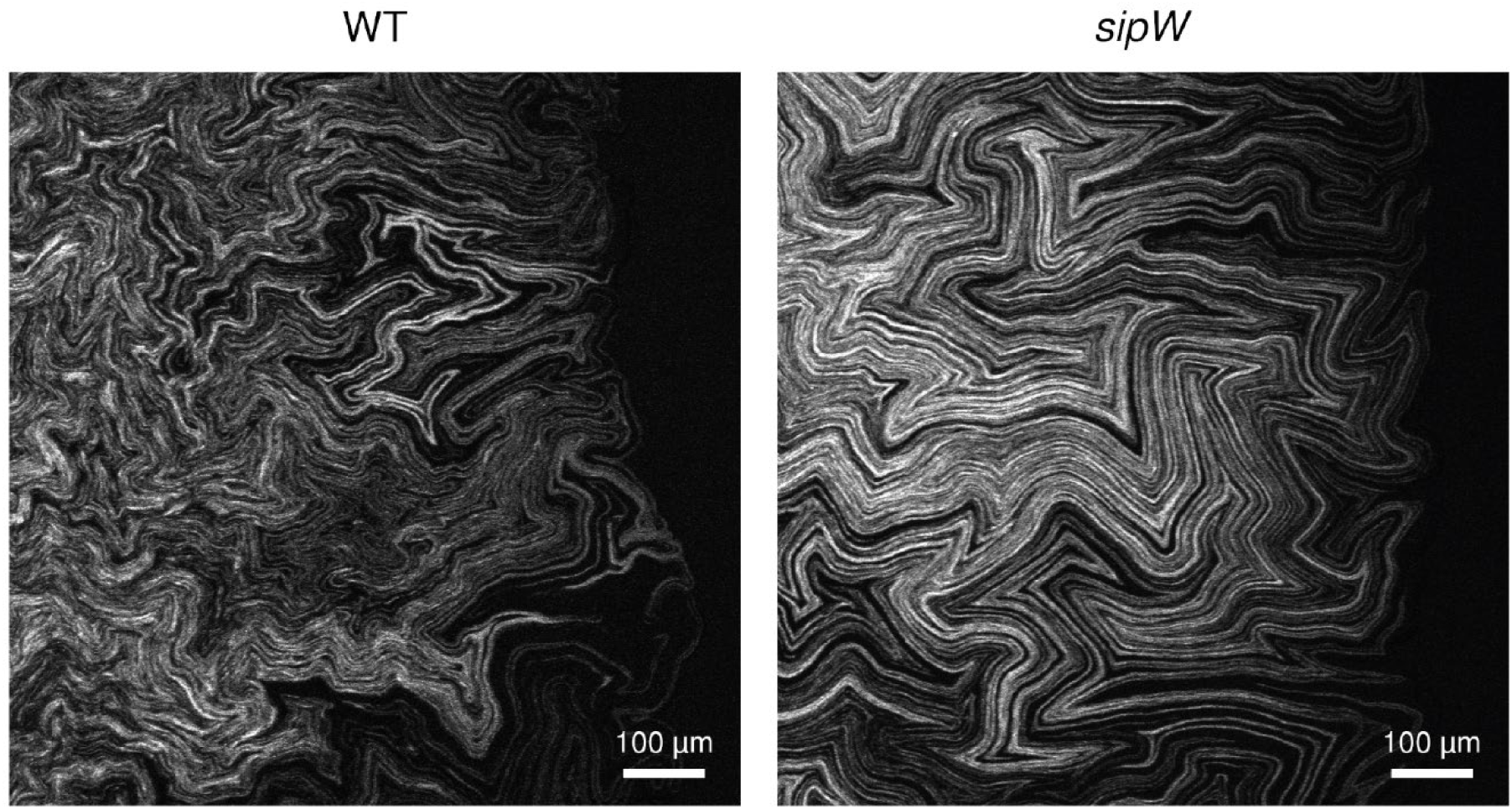
Aerial structure formation at the colony biofilm periphery. Confocal imaging of NCIB 3610 colony biofilm edges show development of complex structures within the width of a field of view, which contrasts with the maintenance of monolayered chains in a sipW colony biofilm. Still images are taken from the time-lapse microscopy of Figure Movie 2 and Figure Movie S2 B respectively. Colony biofilms were formed upon coculture of NCIB 3610 with NRS1473 (WT) and NRS5488 with NRS6718 (*sipW*). Images shown are from approximately 14 hours after cell deposition.

**Figure S 4.**
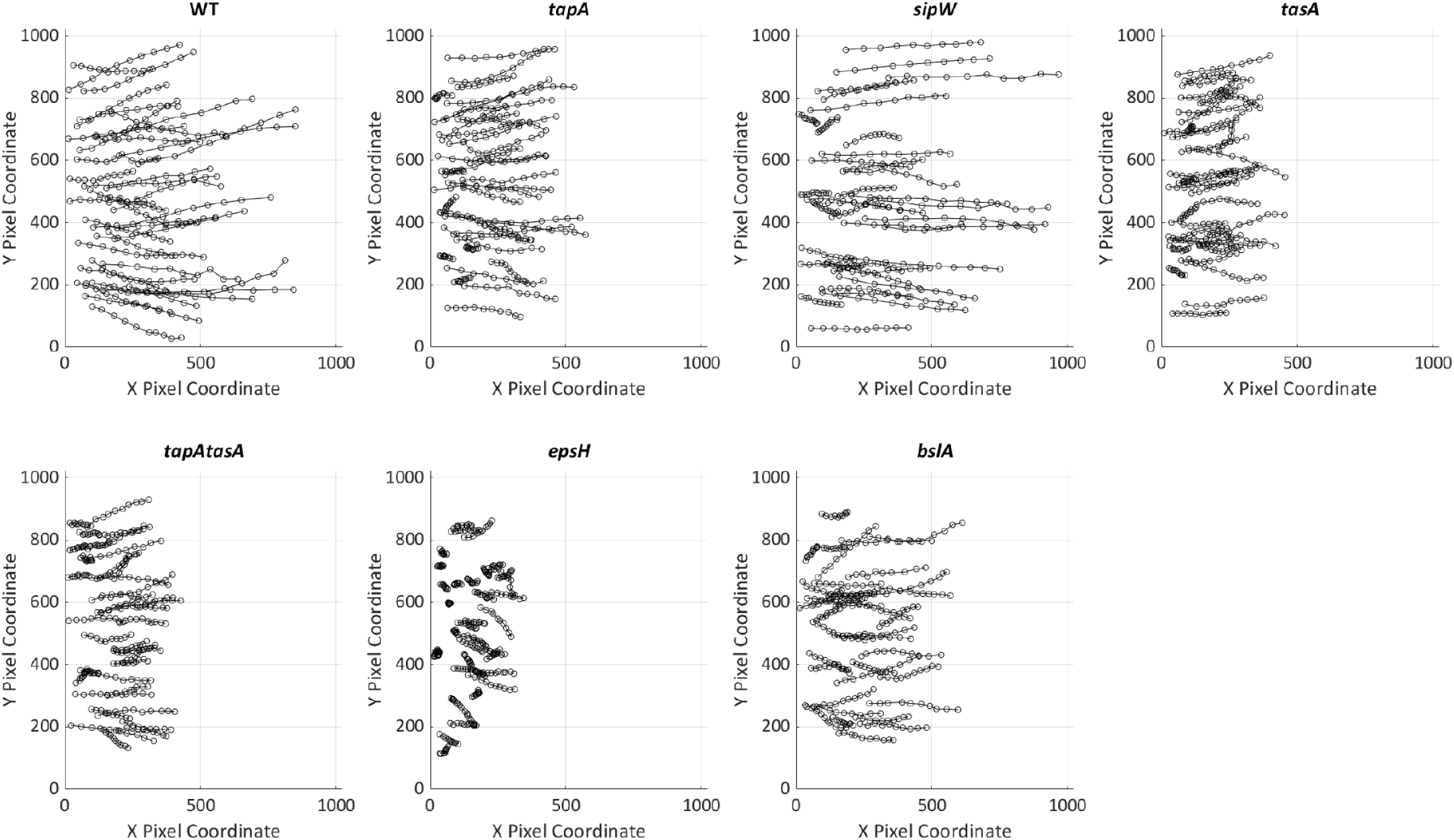
Tracking biofilm landmark movement over time. Visibly recognisable landmarks had their locations specified by placement of a region of interest (ROI) on each time-point in time-lapse imaging (see Fig Movie S3). Tracks are represented by a temporal projection of the coordinates of the ROIs on the 1024 x 1024-pixel image space. In total, 6 tracks from each of 6 time-lapse images are shown per genotype.

**Figure S 5.**
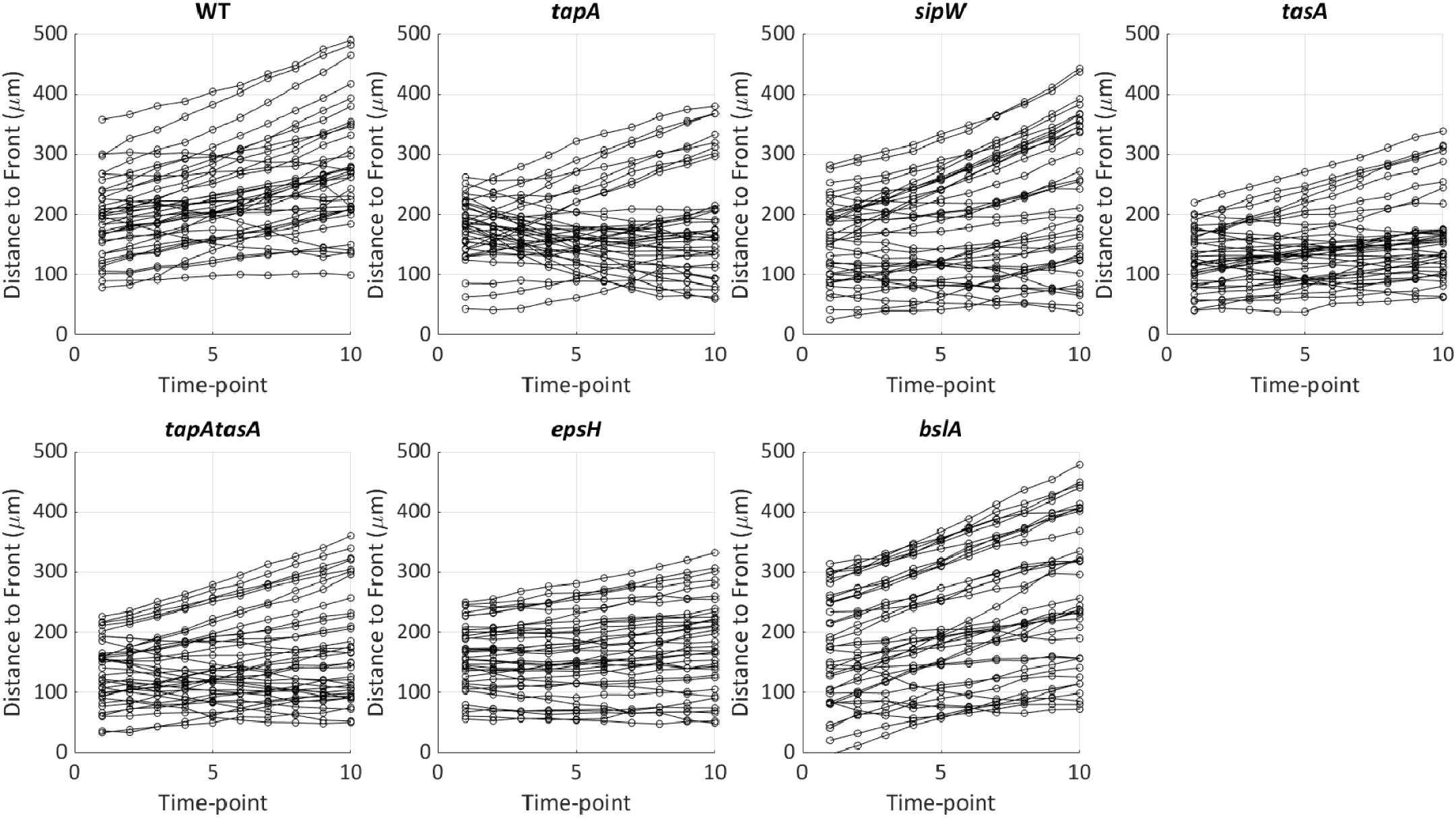
Landmark motion relative to the edge of the biofilm. For each landmark tracked across time-lapse images, the distance to the expanding edge (Distance to Front) of the biofilm was measured at each time-point (see Figure Movie S3, ‘d’). In the plots, approximately horizontal lines show landmarks keeping pace with the expanding edge, whereas positive or negative gradient lines show landmarks falling behind or catching up with the edge, respectively

**Figure Movie S 1.**
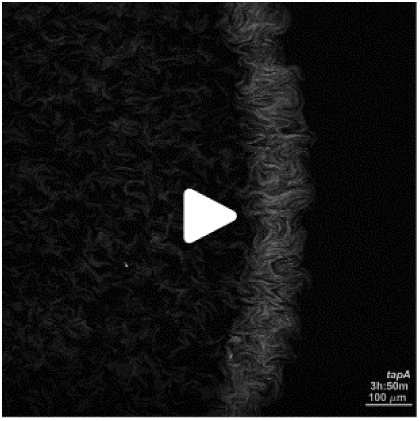
A-F Initial cell growth. Time-lapse confocal microscopy of cocultured strains of the genotypes indicated below containing 20% GFP-producing cells at the point of inoculation from 30 minutes after cell deposition. Z-stacks were acquired every 10 minutes for 10 hours, and a maximum intensity projection is presented. The timestamp is the approximate age of the biofilm since cells were deposited and the scale bar is 100 µm. The strains cocultured were **(A)** *tapA* NRS3936/NRS6723; **(B)** *sipW* NRS5488/NRS6718; **(C)** *tasA* NRS5267/NRS6724; **(D)** *tapA tasA* NRS5748/NRS6727; **(E)** *epsH* NRS5906/NRS6728; **(F)** *bslA* NRS2097/NRS5131).

**Figure Movie S 2.**
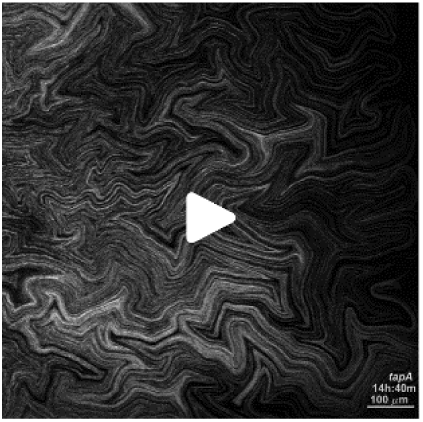
A-F Cell chains emanate from the biofilm periphery. Time-lapse confocal microscopy of cocultured strains of the genotypes indicated below containing 20% GFP-producing cells at the point of inoculation from 10.5 hours after cell deposition and incubation at 30°C. Z-stacks were acquired every 10 minutes for 14 hours, and a maximum intensity projection is presented. The timestamp is the approximate age of the biofilm since cells were deposited and the scale bar is 100 µm. The strains cocultured were **(A)** *tapA* NRS3936/NRS6723; **(B)** *sipW* NRS5488/NRS6718; **(C)** *tasA* NRS5267/NRS6724; **(D)** *tapA tasA* NRS5748/NRS6727; **(E)** *epsH* NRS5906/NRS6728; **(F)** *bslA* NRS2097/NRS5131).

**Figure Movie S 3.**
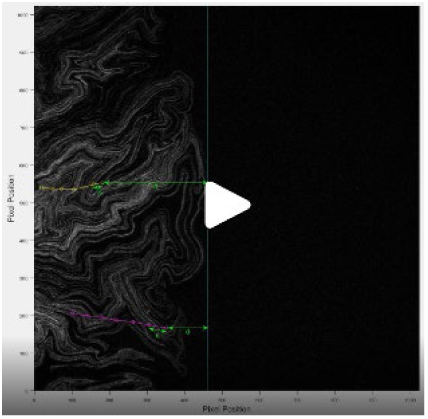
Tracking and measuring landmark motion. Time-lapse confocal microscopy of cocultured strains NCIB 3610 (80%) and GFP-producing NRS1473 (20%) from 10.5 hours after cell deposition and incubation at 30°C. Z-stacks were acquired every 10 minutes for 14 hours, and a maximum intensity projection is presented. In this example, two visibly recognisable landmarks have their locations defined by placement of an ROI at each time-point, represented by the emerging yellow and magenta tracks. Two measurements are then made: step-size ‘s’ is the Euclidean distance a landmark travelled between discrete time-points; measurement ‘d’ is the distance from the landmark to the expanding edge of the biofilm.

**Figure Movie S 4.**
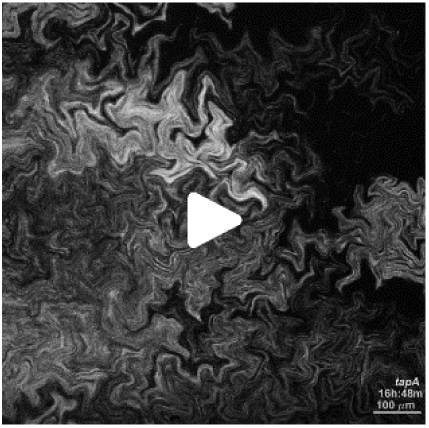
A-F Biomass structure development at the biofilm-agar interface. Time-lapse confocal microscopy at the biofilm-agar interface of mixed strains of the genotypes stated containing 20% GFP-producing cells at the point of inoculation. Imaging started at 12 hours after cell deposition and incubation was at 30°C. Z-stacks were acquired every 18 minutes, and a maximum intensity projection is presented. The timestamp is the approximate age of the biofilm since cells were deposited and the scale bar is 100 µm. The strains cocultured were **(A)** *tapA* NRS3936/NRS6723; **(B)** *sipW* NRS5488/NRS6718; **(C)** *tasA* NRS5267/NRS6724; **(D)** *tapA tasA* NRS5748/NRS6727; **(E)** *epsH* NRS5906/NRS6728; **(F)** *bslA* NRS2097/NRS5131)

**Figure Movie S 5.**
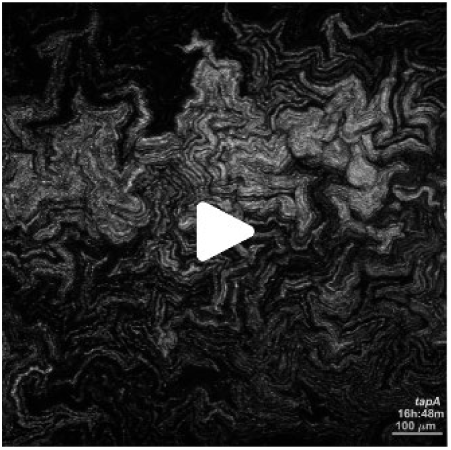
A-F Biomass movement at the biofilm-air interface. Time-lapse confocal microscopy at the biofilm-air interface of mixed strains of the genotypes stated containing 20% GFP-producing cells at the point of inoculation. Imaging started at 12 hours after cell deposition and incubation at 30°C. Z-stacks were acquired every 18 minutes, and a maximum intensity projection is presented. The timestamp is the approximate age of the biofilm since cells were deposited and the scale bar is 100 µm. The strains cocultured were **(A)** *tapA* NRS3936/NRS6723; **(B)** *sipW* NRS5488/NRS6718; **(C)** *tasA* NRS5267/NRS6724; **(D)** *tapA tasA* NRS5748/NRS6727; **(E)** *epsH* NRS5906/NRS6728; **(F)** *bslA* NRS2097/NRS5131)

## References

Allan, C., J.-M. Burel, J. Moore, C. Blackburn, M. Linkert, S. Loynton, D. MacDonald, W.J. Moore, C. Neves, A. Patterson, M. Porter, A. Tarkowska, B. Loranger, J. Avondo, I. Lagerstedt, L. Lianas, S. Leo, K. Hands, R.T. Hay, A. Patwardhan, C. Best, G.J. Kleywegt, G. Zanetti, and J.R. Swedlow. 2012. OMERO: flexible, model-driven data management for experimental biology. Nat. Methods. 9:245–253. doi:10.1038/nmeth.1896.

Arnaouteli, S., N.C. Bamford, N.R. Stanley-Wall, and Á.T. Kovács. 2021. *Bacillus subtilis* biofilm formation and social interactions. Nat. Rev. Microbiol. doi:10.1038/s41579-021-00540-9.

Arnaouteli, S., D.A. Matoz-Fernandez, M. Porter, M. Kalamara, J. Abbott, C.E. MacPhee, F.A. Davidson, and N.R. Stanley-Wall. 2019. Pulcherrimin formation controls growth arrest of the *Bacillus subtilis* biofilm. Proc. Natl. Acad. Sci. U. S. A. 116:13553–13562. doi:10.1073/pnas.1903982116.

Arnaud, M., A. Chastanet, and M. Débarbouillé. 2004. New Vector for Efficient Allelic Replacement in Naturally Nontransformable, Low-GC-Content, Gram-Positive Bacteria. Appl. Environ. Microbiol. 70:6887–6891. doi:10.1128/AEM.70.11.6887-6891.2004.

Branda, S.S., J.E. Gonzalez-Pastor, S. Ben-Yehuda, R. Losick, and R. Kolter. 2001. Fruiting body formation by *Bacillus subtilis*. Proc. Natl. Acad. Sci. 98:11621–11626. doi:10.1073/pnas.191384198.

Cámara-Almirón, J., Y. Navarro, L. Díaz-Martínez, M.C. Magno-Pérez-Bryan, C. Molina-Santiago, J.R. Pearson, A. de Vicente, A. Pérez-García, and D. Romero. 2020. Dual functionality of the amyloid protein TasA in *Bacillus* physiology and fitness on the phylloplane. Nat. Commun. 11:1859. doi:10.1038/s41467-020-15758-z.

Dar, D., N. Dar, L. Cai, and D.K. Newman. 2021. Spatial transcriptomics of planktonic and sessile bacterial populations at single-cell resolution. Science (80-. ). 373. doi:10.1126/science.abi4882.

Dragoš, A., and Á.T. Kovács. 2017. The Peculiar Functions of the Bacterial Extracellular Matrix. Trends Microbiol. 25:257–266. doi:10.1016/j.tim.2016.12.010.

Dragoš, A., Á.T. Kovács, and D. Claessen. 2017. The Role of Functional Amyloids in Multicellular Growth and Development of Gram-Positive Bacteria. Biomolecules. 7:60. doi:10.3390/biom7030060.

Earl, C., S. Arnaouteli, N.C. Bamford, M. Porter, T. Sukhodub, C.E. MacPhee, and N.R. Stanley-Wall. 2020. The majority of the matrix protein TapA is dispensable for *Bacillus subtilis* colony biofilm architecture. Mol. Microbiol. mmi.14559. doi:10.1111/mmi.14559.

Erskine, E., R.J. Morris, M. Schor, C. Earl, R.M.C. Gillespie, K.M. Bromley, T. Sukhodub, L. Clark, P.K. Fyfe, L.C. Serpell, N.R. Stanley-Wall, and C.E. MacPhee. 2018. Formation of functional, non-amyloidogenic fibres by recombinant *Bacillus subtilis* TasA. Mol. Microbiol. 110:897–913. doi:10.1111/mmi.13985.

Fei, C., S. Mao, J. Yan, R. Alert, H.A. Stone, B.L. Bassler, N.S. Wingreen, and A. Košmrlj. 2020. Nonuniform growth and surface friction determine bacterial biofilm morphology on soft substrates. Proc. Natl. Acad. Sci. 117:7622–7632. doi:10.1073/pnas.1919607117.

Flemming, H.C., and J. Wingender. 2010. The biofilm matrix. Nat. Rev. Microbiol. 8:623–633. doi:10.1038/nrmicro2415.

Flemming, H.C., J. Wingender, U. Szewzyk, P. Steinberg, S.A. Rice, and S. Kjelleberg. 2016. Biofilms: An emergent form of bacterial life. Nat. Rev. Microbiol. 14:563–575. doi:10.1038/nrmicro.2016.94.

Gamba, P., J.W. Veening, N.J. Saunders, L.W. Hamoen, and R.A. Daniel. 2009. Two-step assembly dynamics of the *Bacillus subtilis* divisome. J. Bacteriol. 191:4186–4194. doi:10.1128/JB.01758-08.

van Gestel, J., F.J. Weissing, O.P. Kuipers, and Á.T. Kovács. 2014. Density of founder cells affects spatial pattern formation and cooperation in *Bacillus subtilis* biofilms. ISME J. 8:2069–2079. doi:10.1038/ismej.2014.52.

Gordon, V., L. Bakhtiari, and K. Kovach. 2019. From molecules to multispecies ecosystems: the roles of structure in bacterial biofilms. Phys. Biol. 16:041001. doi:10.1088/1478-3975/ab1384.

Hobley, L., A. Ostrowski, F. V Rao, K.M. Bromley, M. Porter, A.R. Prescott, C.E. MacPhee, D.M.F. van Aalten, and N.R. Stanley-Wall. 2013. BslA is a self-assembling bacterial hydrophobin that coats the *Bacillus subtilis* biofilm. Proc. Natl. Acad. Sci. 110:13600–13605. doi:10.1073/pnas.1306390110.

Kesel, S., B. von Bronk, C. Falcón García, A. Götz, O. Lieleg, and M. Opitz. 2017. Matrix composition determines the dimensions of *Bacillus subtilis* NCIB 3610 biofilm colonies grown on LB agar. RSC Adv. 7:31886–31898. doi:10.1039/C7RA05559E.

Krajnc, M., P. Stefanic, R. Kostanjšek, I. Mandic-Mulec, I. Dogsa, and D. Stopar. 2022. Systems view of *Bacillus subtilis* pellicle development. npj Biofilms Microbiomes. 8:1–11. doi:10.1038/s41522-022-00293-0.

Meacock, O.J., A. Doostmohammadi, K.R. Foster, J.M. Yeomans, and W.M. Durham. 2020. Bacteria solve the problem of crowding by moving slowly. Nat. Phys. doi:10.1038/s41567-020-01070-6.

O’Toole, G., H.B. Kaplan, and R. Kolter. 2000. Biofilm Formation as Microbial Development. Annu. Rev. Microbiol. 54:49–79. doi:10.1146/annurev.micro.54.1.49.

O’Toole, G.A., and R. Kolter. 1998. Flagellar and twitching motility are necessary for Pseudomonas aeruginosa biofilm development. Mol. Microbiol. 30:295–304. doi:10.1046/J.1365-2958.1998.01062.X.

Ostrowski, A., A. Mehert, A. Prescott, T.B. Kiley, and N.R. Stanley-Wall. 2011. YuaB Functions Synergistically with the Exopolysaccharide and TasA Amyloid Fibers To Allow Biofilm Formation by *Bacillus subtilis*. J. Bacteriol. 193:4821–4831. doi:10.1128/JB.00223-11.

Otsu, N. 1979. A Threshold Selection Method from Gray-Level Histograms. IEEE Trans. Syst. Man. Cybern. 9:62–66. doi:10.1109/TSMC.1979.4310076.

Romero, D., C. Aguilar, R. Losick, and R. Kolter. 2010. Amyloid fibers provide structural integrity to *Bacillus subtilis* biofilms. Proc. Natl. Acad. Sci. 107:2230–2234. doi:10.1073/pnas.0910560107.

Rooney, L.M., W.B. Amos, P.A. Hoskisson, and G. McConnell. 2020. Intra-colony channels in *E. coli* function as a nutrient uptake system. ISME J. doi:10.1038/s41396-020-0700-9.

Roux, D., C. Cywes-Bentley, Y.F. Zhang, S. Pons, M. Konkol, D.B. Kearns, D.J. Little, P.L. Howell, D. Skurnik, and G.B. Pier. 2015. Identification of Poly-N-acetylglucosamine as a major polysaccharide component of the *Bacillus subtilis* biofilm matrix. J. Biol. Chem. 290:19261–19272. doi:10.1074/jbc.M115.648709.

Sanchez-Vizuete, P., Y. Dergham, A. Bridier, J. Deschamps, E. Dervyn, K. Hamze, S. Aymerich, D. Le Coq, and R. Briandet. 2022. The coordinated population redistribution between *Bacillus subtilis* submerged biofilm and liquid-air pellicle. Biofilm. 4:100065. doi:10.1016/j.bioflm.2021.100065.

Sarkans, U., M. Gostev, A. Athar, E. Behrangi, O. Melnichuk, A. Ali, J. Minguet, J.C. Rada, C. Snow, A. Tikhonov, A. Brazma, and J. McEntyre. 2018. The BioStudies database—one stop shop for all data supporting a life sciences study. Nucleic Acids Res. 46:D1266–D1270. doi:10.1093/nar/gkx965.

Seminara, A., T.E. Angelini, J.N. Wilking, H. Vlamakis, S. Ebrahim, R. Kolter, D.A. Weitz, and M.P. Brenner. 2012. Osmotic spreading of *Bacillus subtilis* biofilms driven by an extracellular matrix. Proc. Natl. Acad. Sci. 109:1116–1121. doi:10.1073/pnas.1109261108.

Srinivasan, S., I.D. Vladescu, S.A. Koehler, X. Wang, M. Mani, and S.M. Rubinstein. 2018. Matrix Production and Sporulation in *Bacillus subtilis* Biofilms Localize to Propagating Wave Fronts. Biophys. J. 114:1490–1498. doi:10.1016/j.bpj.2018.02.002.

Stanley, N.R., R.A. Britton, A.D. Grossman, and B.A. Lazazzera. 2003. Identification of Catabolite Repression as a Physiological Regulator of Biofilm Formation by *Bacillus subtilis* by Use of DNA Microarrays. J. Bacteriol. 185:1951–1957. doi:10.1128/JB.185.6.1951-1957.2003.

Terra, R., N.R. Stanley-Wall, G. Cao, and B.A. Lazazzera. 2012. Identification of *Bacillus subtilis* SipW as a Bifunctional Signal Peptidase That Controls Surface-Adhered Biofilm Formation. J. Bacteriol. 194:2781–2790. doi:10.1128/JB.06780-11.

Tjalsma, H., A. Bolhuis, J.D.H. Jongbloed, S. Bron, and J.M. van Dijl. 2000. Signal Peptide-Dependent Protein Transport in*Bacillus subtilis*: a Genome-Based Survey of the Secretome. Microbiol. Mol. Biol. Rev. 64:515–547. doi:10.1128/MMBR.64.3.515-547.2000.

Veening, J.-W., W.K. Smits, L.W. Hamoen, and O.P. Kuipers. 2006. Single cell analysis of gene expression patterns of competence development and initiation of sporulation in *Bacillus subtilis* grown on chemically defined media. J. Appl. Microbiol. 101:531–541. doi:10.1111/j.1365-2672.2006.02911.x.

Verhamme, D.T., T.B. Kiley, and N.R. Stanley-Wall. 2007. DegU co-ordinates multicellular behaviour exhibited by *Bacillus subtilis*. Mol. Microbiol. 65:554–568. doi:10.1111/j.1365-2958.2007.05810.x.

Vlamakis, H., C. Aguilar, R. Losick, and R. Kolter. 2008. Control of cell fate by the formation of an architecturally complex bacterial community. Genes Dev. 22:945–953. doi:10.1101/gad.1645008.

Wang, X., S.A. Koehler, J.N. Wilking, N.N. Sinha, M.T. Cabeen, S. Srinivasan, A. Seminara, S. Rubinstein, Q. Sun, M.P. Brenner, and D.A. Weitz. 2016. Probing phenotypic growth in expanding *Bacillus subtilis* biofilms. Appl. Microbiol. Biotechnol. 100:4607–4615. doi:10.1007/s00253-016-7461-4.

Warren, M.R., H. Sun, Y. Yan, J. Cremer, B. Li, and T. Hwa. 2019. Spatiotemporal establishment of dense bacterial colonies growing on hard agar. Elife. 8:1–47. doi:10.7554/eLife.41093.

Watnick, P.I., C.M. Lauriano, K.E. Klose, L. Croal, and R. Kolter. 2001. The absence of a flagellum leads to altered colony morphology, biofilm development and virulence in *Vibrio cholerae* O139. Mol. Microbiol. 39:223–235. doi:10.1046/j.1365-2958.2001.02195.x.

Yaman, Y.I., E. Demir, R. Vetter, and A. Kocabas. 2019. Emergence of active nematics in chaining bacterial biofilms. Nat. Commun. 10:2285. doi:10.1038/s41467-019-10311-z.

Yan, J., A.G. Sharo, H.A. Stone, N.S. Wingreen, and B.L. Bassler. 2016. *Vibrio cholerae* biofilm growth program and architecture revealed by single-cell live imaging. Proc. Natl. Acad. Sci. 113:e5337–e5343. doi:10.1073/pnas.1611494113.

Zhang, W., W. Dai, S.-M. Tsai, S.M. Zehnder, M. Sarntinoranont, and T.E. Angelini. 2015. Surface indentation and fluid intake generated by the polymer matrix of *Bacillus subtilis* biofilms. Soft Matter. 11:3612–3617. doi:10.1039/C5SM00148J.

